# Mucoricin is a Ricin-Like Toxin that is Critical for the Pathogenesis of Mucormycosis

**DOI:** 10.1101/2021.01.13.426274

**Authors:** Sameh S. M. Soliman, Clara Baldin, Yiyou Gu, Shakti Singh, Teclegiorgis Gebremariam, Marc Swidergall, Abdullah Alqarihi, Eman G. Youssef, Sondus Alkhazraji, Antonis Pikoulas, Christina Perske, Vivek Venkataramani, Abigail Rich, Vincent M. Bruno, Julie Dunning Hotopp, Nicolas J. Mantis, John E. Edwards, Scott G. Filler, Georgios Chamilos, Ellen S. Vitetta, Ashraf S. Ibrahim

**Affiliations:** Division of Infectious Diseases, The Lundquist Institute for Biomedical Innovation at Harbor-University of California at Los Angeles (UCLA) Medical Center, Torrance, CA, U.S.A.; College of Pharmacy and Research Institute for Medical and Health Sciences, University of Sharjah, P.O. Box 27272, Sharjah, United Arab Emirates; Department of Biological Chemistry, Medical University of Innsbruck, Innsbruck, Austria; David Geffen School of Medicine at UCLA, Los Angeles, Ca, U.S.A; Department of Biotechnology and Life Sciences, Faculty of Postgraduate Studies for Advanced Sciences, Beni-Suef University, Beni-Suef, Egypt; Department of Medicine, University of Crete, Foundation for Research and Technology, 71300 Heraklion, Crete, Greece; Department of Pathology, University Medicine Göttingen, Göttingen, Germany; Department of Hematology and Medical Oncology, University Medicine Göttingen, Göttingen, Germany; Departments of Immunology and Microbiology, UT Southwestern Medical Center, Dallas, Tx, U.S.A; Department of Microbiology and Immunology, University of Maryland School of Medicine, Baltimore, MD, U.S.A; New York State Department of Health, Wadsworth Center, Albany, NY, U.S.A.

**Keywords:** Mucormycosis, Toxins, Ricin, *Rhizopus*, Virulence, Pathogenesis, Mucoricin

## Abstract

Fungi of the order Mucorales cause mucormycosis, a lethal infection with an incompletely understood pathogenesis. We now demonstrate that Mucorales fungi produce a toxin that plays a central role in virulence. Polyclonal antibodies against this toxin inhibit its ability to damage human cells *in vitro,* and prevent hypovolemic shock, organ necrosis, and death in mice with mucormycosis. RNAi inhibition of the toxin in *Rhizopus delemar,* compromises the ability of the fungus to damage host cells and attenuates virulence in mice. This 17 kDa toxin has structural and functional features of the plant toxin, ricin, including the ability to inhibit protein synthesis by its N-glycosylase activity, the existence of a motif that mediates vascular leak, and a lectin sequence. Antibodies against the toxin inhibit *R. delemar-* or toxin-mediated vascular permeability *in vitro* and cross-react with ricin. A monoclonal anti-ricin B chain antibody binds to the toxin and also inhibits its ability to cause vascular permeability. Therefore, we propose the name “mucoricin” for this toxin. Not only is mucoricin important in the pathogenesis of mucormycosis but our data suggest that a ricin-like toxin is produced by organisms beyond the plant and bacterial kingdoms. Importantly, mucoricin should be a promising therapeutic target.

---

Mucormycosis is a lethal fungal infection that usually afflicts immunocompromised hosts such as diabetics in ketoacidosis (DKA), neutropenic patients, patients undergoing hematopoietic cell or solid organ transplant, or patients receiving high-dose corticosteroids^1–6^. Immunocompetent patients with severe trauma are also at risk of contracting mucormycosis by direct inoculation of open wounds^7,8^. The overall mortality rate of mucormycosis is >40% and it approaches 100% in patients with disseminated disease, persistent neutropenia, or brain infection^1–6^. The two most common forms of the disease are rhino-orbital/cerebral and pulmonary mucormycosis. In both forms of the disease, infection is initiated by the inhalation of spores that germinate in the host to form hyphae, which are capable of invading host tissues while avoiding phagocytic killing^6,9^.

A characteristic feature of mucormycosis is the propensity of Mucorales to invade blood vessels, resulting in thrombosis and subsequent tissue necrosis^6^. The massive tissue necrosis associated with mucormycosis compromises the delivery of antifungal drugs to infected foci, thereby necessitating radical surgical intervention to improve the outcome of therapy. We have previously determined that Mucorales fungi invade human umbilical vein endothelial cells (HUVECs) by expressing the fungal invasin, CotH3, which interacts with the 78kDa host receptor, <u>g</u>lucose <u>r</u>egulated <u>p</u>rotein (GRP78). The interaction between CotH3 and GRP78 induces the endothelial cells to endocytose the fungi^10–12^. However, the mechanisms by which Mucorales damage host cells and cause necrosis are unknown.

While studying the capacity of *Rhizopus delemar,* the most common cause of mucormycosis, to damage HUVECs, we observed that killed hyphae of this organism and other Mucorales caused considerable damage to host cells^13^. This experimental finding and the clinical observation of the extensive tissue necrosis observed in patients with mucormycosis led us to speculate that a fungal-derived toxin may be involved in the pathogenesis of this disease.

Here, we identify and characterize a hyphal-associated and secreted/shed toxin produced by Mucorales. This toxin damages host cells *in vitro* by inhibiting protein synthesis. The toxin is required for the pathogenesis of mucormycosis in mice, where it induces inflammation, hemorrhage and tissue damage resulting in apoptosis and necrosis. Suppression of toxin production in *R. delemar* by RNAi attenuates virulence in DKA mice, and polyclonal anti-toxin antibodies (IgG anti-toxin) protect mice from mucormycosis by reducing tissue inflammation and damage. Thus, the toxin is a key virulence factor of Mucorales fungi and a promising therapeutic target. Because this toxin shares structural and functional features with ricin produced by the castor bean plant, *Ricinus communis*^14^, we named it “mucoricin”.

## Results

### Mucorales damage host cells by a hyphal-associated toxin

We previously observed that *R. delemar* causes significant damage to HUVECs within 8 h of infection^11^. This organism also damages the A549 alveolar epithelial cell line and primary alveolar epithelial cells, but only after 30 h of incubation (**Extended Data Fig. 1a**). *R. delemar*-mediated damage to both HUVECs and alveolar epithelial cells is associated with the formation of extensive hyphae, suggesting that the hyphal form of this organism produces a factor(s) that damage host cells^13^. To investigate whether viability is required for *R. delemar* hyphae to damage host cells, we compared the extent of damage to A549 cells caused by live and heat-killed hyphae. We found that while heat-killed hyphae caused less damage to these cells than live hyphae, the extent of host cell damage was still significant (**Extended Data Fig. 1b)**. These finding suggested that a hyphal-associated heat-stable toxin may be partially responsible for host cell damage. To explore this hypothesis, we compared the ability of aqueous extracts from dead *R. delemar* spores and/or hyphae to damage host cells. Extracts from either hyphae alone or from a mixture of spores and hyphae damaged A549 cells, whereas an extract from spores alone caused no detectable damage (**Extended Data Fig. 1c**). We also found that killed cells and pelleted hyphal debris from four different Mucorales fungi, but not the yeast *Candida albicans,* caused significant damage to HUVECs (**Extended Data Fig. 1d**). Collectively, these results suggest that Mucorales produce a hyphal-associated toxin that damages mammalian cells.

### Purification and activity of the toxin

To purify the hyphal-associated toxin, *R. delemar* spores were grown in a liquid medium for 4-7 days to generate a hyphal mat. The mat was ground in liquid nitrogen and extracted with sterile water, concentrated and analyzed by size exclusion chromatography^15^. When the different fractions were analyzed for their ability to damage A549 cells, activity was found in the fractions with molecular masses of 10-30 kDa (**Extended Data Fig. 2a**). The concentrated water extract was then subjected to three dimensional chromatographic fractionations yielding a fraction that caused significant damage to A549 cells (**Extended Data Fig. 2b-g**). This fraction was further subjected to high-performance liquid chromatography (HPLC), using hydrophobic interaction chromatography (HIC). The purified sample was trypsinized and sequenced using LC-MS/MS, which identified a 17 kDa protein (RO3G_06568).

The ORF encoding the 17 kDa protein is widely present in other Mucorales that we and others previously sequenced^16–19^ and that are reported to cause disease in humans (*Mucor, Cunninghamella, Lichtheimia*), animals (*Mortierella*), or plants (*Choanephora cucurbitarum*). Orthologues were also found in the arbuscular mycorrhizal fungus *Rhizophagus* species, and bacterial genera of *Streptomyces* and *Paenibacillus* (**Supplementary Table 1**). Since orthologues were detected in other Mucorales known to cause mucormycosis, we examined the ability of unfractionated hyphal extracts from various Mucorales fungi to damage A549 cells relative to that induced by hyphal extracts from the *R. delemar* 99-880 reference strain. Hyphal extracts from *R. oryzae*, another strain of *R. delemar*, *Lichtheimia corymbifera*, and *Cunninghamella bertholletiae* all caused significant damage to A549 cells (**Extended Data Fig. 1e**).

We expressed the 17 kDa putative toxin gene in *S. cerevisiae* and used the purified recombinant toxin to raise polyclonal anti-toxin antibodies in rabbits. Although the IgG fraction of the antisera (IgG anti-toxin) had no effect on the growth or germination of *R. delemar in vitro* (**Extended Data Fig. 3**), it resulted in ∼50%-70% inhibition of the damage to A549 cells caused by heat-killed hyphae of several Mucorales (**Extended Data Fig. 1f**). From these findings we concluded that the putative toxin is responsible for host cell damage caused by most, if not all, members of the Mucorales fungi.

We also used qRT-PCR to study the expression of this ORF in *R. delemar.* In accordance with data showing a lack of toxin activity in spores (**Extended Data Fig. 1c**), there was minimal expression of this gene during the first 3 h of incubation (prior to germination^13^). Expression of this gene began to increase when the spores germinated at 4 h^13^, peaked by 5 h, and plateaued for at least 16 h of hyphal formation (**Extended Data Fig. 4a**). Protein expression in germlings and hyphae, but not spores, was confirmed by immunostaining using the IgG anti-toxin (**Extended Data Fig. 4b**). The expression of the putative toxin gene was high in hyphae growing under aerated conditions, but not in hyphae grown in the absence of aeration (**Extended Data Fig. 4c).** In addition, RNA expression was 5-10-fold higher following 2-5 h of co-culture with A549 alveolar epithelial cells as compared to co-culture with HUVECs or human erythrocytes. (**Extended Data Fig. 4d**).

### The toxin is capable of damaging host cells *in vitro* and *in vivo*

We compared the ability of the purified toxin to damage primary lung epithelial cells, A549 alveolar epithelial cells, and HUVECs. After 1 h, the toxin caused significant damage to all the host cells and especially to HUVECs (**Fig. 1a**). After 3 h, there was almost 100% damage to all host cells. We also compared the ability of the purified toxin *vs.* the recombinant toxin to damage A549 cells. Both caused significant damage (**Fig. 1b**). Therefore, the purified and the recombinant protein act similarly and in a time dependent manner in damaging A549 cells.

**Figure 1.**
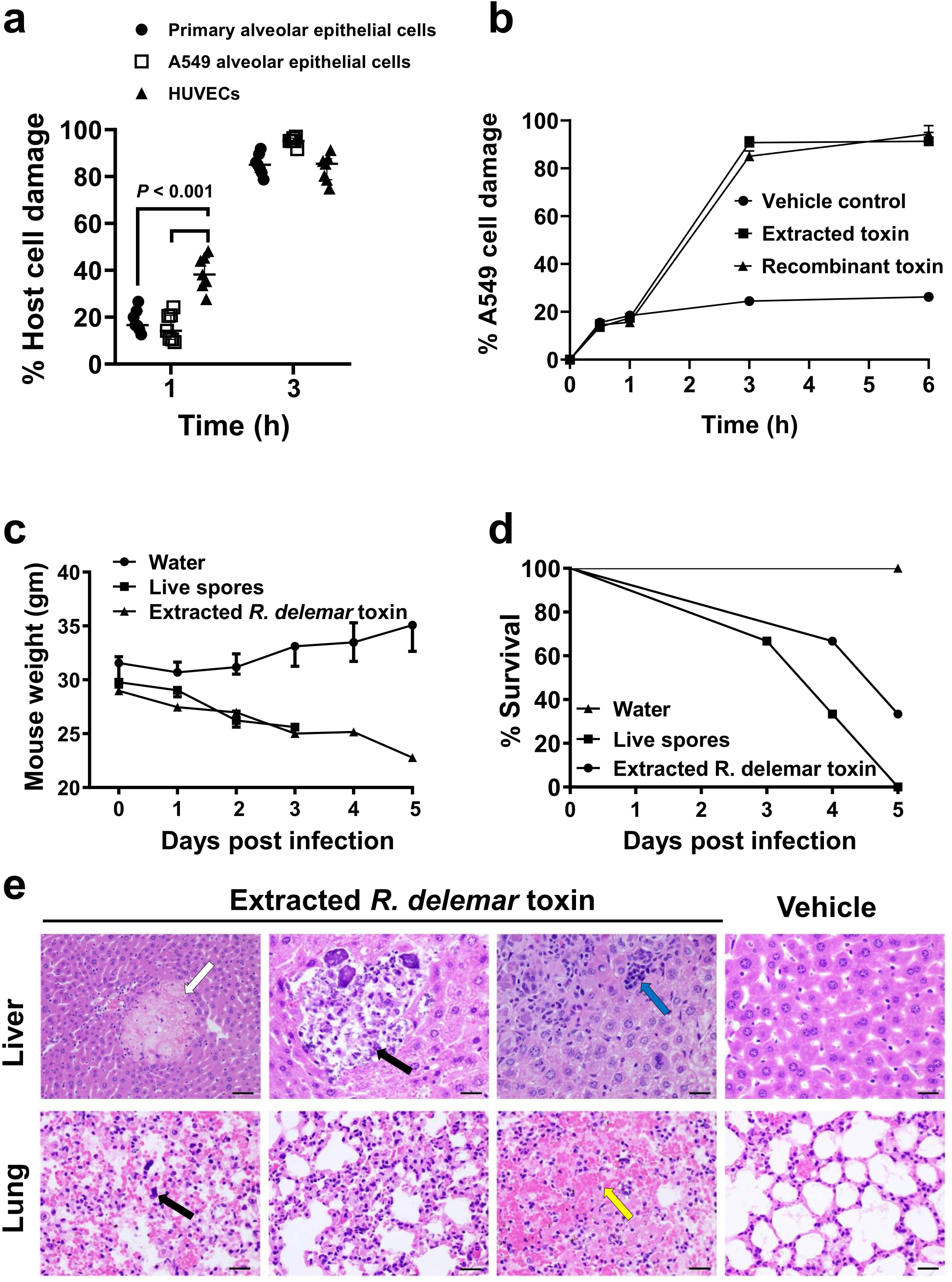
*R. delemar* toxin is sufficient to cause damage *in vitro* and *in vivo.* (**a**) The effect of toxin on different cell lines (n=7 wells /time point pooled from three independent experiments). Data are median ± interquartile range. Statistical analysis was performed by using the non-parametric Mann-Whitney (two-tailed) test comparing HUVECs *vs.* primary alveolar epithelial cells or A549 alveolar cells. **(b)** Damage of extracted or recombinant toxin (∼ 500 μg/ml or 29.4 μM) on epithelial cells at different time points (n=3 wells/time point). Data are representative of three independent experiments and presented as median ± interquartile range. (**c**) Mouse (n=3 mice/group) weight loss (data are median ± interquartile range) and **(d**) percent survival (n=3 mice/group) following intravenous injection with 0.1 mg/ml (5.9 μM) toxin QOD x 3. **(e**) Mouse organ H&E histomicrographs showing the effects of the toxin. Livers showed necrosis (white arrow), infiltration and calcification of PMNs (black arrow) due to inflammation and a cluster of mononuclear cells (cyan arrow). Lungs showed megakaryocytes (black arrow) and hemorrhage (yellow arrow). Data in each group are representative of 2 mice. Scale bar 50 µm for first liver micrograph and 100 µm for all other. For lung micrographs scale bar 50 µm.

We next tested the activity of the toxin *in vivo*. Toxin purified from *R. delemar* was injected intravenously into mice every other day for a total of three doses and the mice were monitored for behavioral changes, weight loss, and survival. Within 10-30 minutes after the injection of 0.1 mg/ml (5.9 μM) purified toxin, we observed behavior highly suggestive of sudden circulatory hypovolemic shock, including rapid and shallow breathing, weakness, and cold skin. The mice lost >25% of their original body weight (**Fig. 1c**); most eventually died. These events were similar to those observed in mice infected with live *R. delemar* spores (**Fig. 1d**). Finally, histopathology of organs collected from the mice showed pathological changes that included necrosis, hemorrhage, and infiltration of the pulmonary interstitium by macrophages in the lungs. Liver changes included necrosis, clusters of mononuclear cells and the presence of megakaryocytes in the organs, polymorphonuclear cell (PMN) infiltration and tissue calcification indicative of uncontrolled inflammation, hemorrhage and necrosis (**Fig. 1e**). These data suggest that the toxin is sufficient to cause clinical symptoms often associated with disseminated mucormycosis.

### RNAi knockdown and antibody-mediated neutralization of the toxin reduced the virulence of *R. delemar in vitro* and *in vivo*

To further confirm the role of this toxin in the pathogenesis of mucormycosis, we used RNAi^20^ to down regulate the gene expression of the toxin. The extent of down regulation of the toxin was measured by qRT-PCR using toxin specific primers and by Western blotting or immunostaining of *R. delemar* with the IgG anti-toxin. This IgG anti-toxin specifically recognized the toxin by ELISA and Western blotting. RNAi knockdown of the toxin caused ∼90% inhibition in gene expression (**Extended Data Fig. 5a**). Furthermore, RNAi knockdown resulted in >80% reduction in protein expression (**Extended Data Fig. 5b**) and negligible staining of toxin-RNAi *R. delemar* germlings compared to germlings of a control strain that have been transformed with an empty plasmid (**Extended Data Fig. 5c**). In accord with the lack of an effect by the IgG anti-toxin on the growth and germination of *R. delemar*, the RNAi knockdown of the toxin had no effect on fungal germination or growth (**Extended Data Fig. 6**).

We next assessed the effect of downregulation of toxin expression on the ability of *R. delemar* to damage A549 alveolar epithelial cells. *R. delemar* with RNAi targeting of the toxin gene induced ∼40% reduction in epithelial cell damage relative to either the wild-type strain or *R. delemar* transformed with the empty plasmid (**Fig. 2a**). Similarly, the IgG anti-toxin protected alveolar epithelial cells from wild-type *R. delemar-*induced injury by ∼40%, *in vitro* whereas normal rabbit IgG did not (**Fig. 2b**).

**Figure 2.**
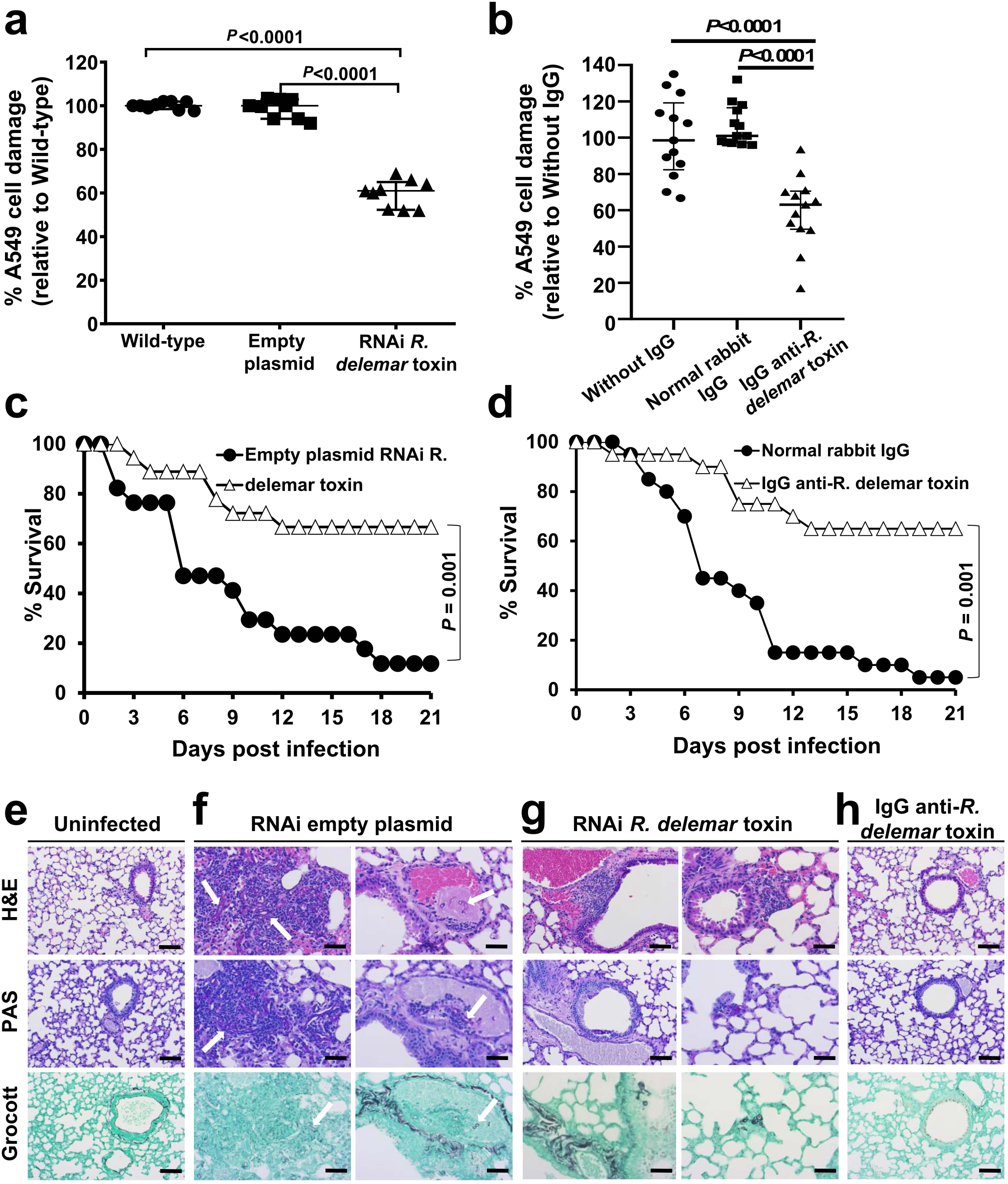
Inhibition of *R. delemar* toxin attenuates virulence of *R. delemar.* **(a)** RNAi toxin shows reduced damage to A549 cells compared to wild type or empty plasmid *R. delamar* (n=6 wells/group pooled from three independent experiments). Data are median ± interquartile range. Statistical comparisons are by the non-parametric Mann-Whitney (two-tailed) test. (**b**) IgG anti-toxin antibodies reduced *R. delemar-*induced injury of A549 cells compared to *R. delamar* without IgG or normal rabbit IgG (n = 13 wells/group pooled from three independent experiments). Data are median ± interquartile range. Statistical comparisons are by the non-parametric Mann-Whitney (two-tailed) test. **(c)** RNAi toxin inhibition prolonged survival of mice (n= 18 mice) compared to *R. delamar* with empty plasmid (n= 17 mice). Data were pooled from two independent experiments. Survival data were analyzed by Log-rank (Mantel-Cox) test. **(d)** IgG anti-toxin prolonged survival of mice compared to normal rabbit IgG (n=20 mice/group). Data were pooled from two independent experiments. Survival data were analyzed by Log-rank (Mantel-Cox) test. **(e)** Histopathological sections of lungs from uninfected mice, **(f)** mice infected with the RNAi empty plasmid *R. delemar* strain showed hyphae and granulocyte infiltration (left panel, arrows) and angioinvasion (right panel, arrow), *vs.* (**g**) mild signs of inflammation and no angioinvasion for mice infected with RNAi toxin. (**h**) IgG anti-toxin group had normal lung tissue architecture. Data in **e-h** are representative of 3 mice and scale bar is 20 µm.

Finally, we evaluated the effects of RNAi inhibition of toxin production on the virulence of *R. delemar* in our model of pulmonary mucormycosis^21^. DKA mice infected with *R. delemar* harboring the empty plasmid had a median survival time of 6 days and 90% mortality by day 21 post-intratracheal infection, whereas mice infected with the toxin-attenuated expression strain had a median survival time of 21 days and mortality of 30% (**Fig. 2c**). Surviving mice had no residual fungal colonies in their lungs when the experiment was terminated on day 21. Inhibition of toxin production appeared to have minimal effects on the early stages of infection because after 4 days of infection, the fungal burden of the lungs and brains (the primary and secondary target organs, respectively^21^) of mice infected with *R. delemar* toxin-attenuated strain and *R. delemar* harboring the empty plasmid were similar (**Extended Data Fig. 7a**). Reduced virulence without affecting tissue fungal burden has been reported for non-neutropenic mice infected with an *Aspergillus fumigatus* null mutant that does not produce the Asp f1 ribotoxin^22^, representing a classical feature of disease tolerance^23^. Collectively, our results indicate that while the toxin is dispensable for the initiation of mucormycosis, it plays a central role in the lethality of this disease.

### IgG anti-toxin protected mice from mucormycosis

To further verify the role of the toxin in the pathogenesis of mucormycosis, we infected DKA mice intratracheally with wild-type *R. delemar* and then 24 h later, treated them with a single 30 µg dose of either the IgG anti-toxin or normal rabbit IgG. While mice treated with normal IgG had a mortality rate of 95%, treatment with the IgG anti-toxin resulted in ∼70% long-term survival (**Fig. 2d**). Surviving mice appeared healthy and had no detectable fungal colonies in their lungs when the experiment was terminated on day 21. In accord with data from the fungal burden in tissues of mice infected with the toxin-attenuated strain, antibody treatment had no effect on the fungal burden of lungs or brains when tissues were harvested four days post infection (**Extended Data Fig. 7b**). These data further confirm the role of the toxin in the pathogenesis of mucormycosis and point to the potential of using anti-toxin antibodies to treat the disease.

We also performed histopathological examination of the tissues from all groups of mice sampled at the same time of tissue fungal burden studies (day 4) to gain insight into the mechanism of action of the toxin. While, uninfected mice had normal lung architecture with no signs of inflammation or infection (**Fig. 2e**), lungs from mice infected with *R. delemar* transformed with the RNAi empty plasmid (control) showed fungal and granulocyte infiltration (**Fig. 2f left panel**) and angioinvasion with thrombosis (**Fig. 2f right panel**). In contrast, lungs from mice infected with the toxin-attenuated mutant showed only mild signs of inflammation with no angioinvasion (**Fig. 2g**). Importantly, lungs of mice infected with wild-type *R. delemar* and treated with the IgG anti-toxin showed architecture that was similar to the lungs of the uninfected control mice; there were no signs of inflammation or infiltration with *R. delemar* (**Fig. 2h**).

### Down regulation of the toxin gene or treatment with IgG anti-toxin attenuated *R. delemar*-mediated host cell damage *in vivo*

To determine whether the toxin contributed to host cell damage *in vivo*, we used the ApopTag *in situ* apoptosis kit to stain the lung tissues of the infected mice. While extensive lung damage was observed in mice infected with wild-type *R. delemar*, the lungs harvested from mice infected with the toxin-attenuated mutant (**Extended Data Fig. 8a**) or those infected with wild-type *R. delemar* and treated with the IgG anti-toxin (**Extended Data Fig. 8b**) had almost no detectable damage.

Finally, we have previously reported on a mucormycosis case in a human with disseminated mucormycosis^9^. Haemotoxylin and Eosin (H&E) staining of lung tissues of this patient showed broad aseptate hyphae that caused necrosis and massive infiltration of tissues compared to thinner septated hyphae present in a patient suffering from invasive pulmonary aspergillosis (**Extended Data Fig. 9a,b**). Subsequent immunohistochemistry of the patient’s lungs using the IgG anti-toxin (*vs.* control IgG) showed association of the toxin with fungal hyphae and the surrounding tissues in a mucormycosis patient and lack of staining in tissues of an aspergillosis patient (**Extended Data Fig. 9c,d**). These results show that the toxin is also involved in human mucormycosis, is cell-associated as well as secreted/shed into the surrounding tissues, and confirm the specificity of the antibody used in these studies since the putative toxin does not have an orthologue in *Aspergillus* (**Supplementary Table 1)**.

To confirm the secretion/shedding of the toxin, we grew *R. delemar* spores in 96-well plate with or without amphotericin B (AmB) and assayed cell-free supernatants for the presence of the toxin using a sandwich ELISA with an IgG_1_ anti-toxin monoclonal antibody that we raised in mice as a capture antibody and the rabbit IgG anti-toxin as the detecting antibody. The toxin was detected in cell-free supernatants of *R. delemar* wild-type (26.7 ± 0.87 nM) or *R. delemar* transformed with the empty plasmid (23.0 ± 2.04 nM), but not in *R. delemar* transformed with RNAi targeting the toxin. In accord with secretion/shedding of the toxin by live hyphae, supernatants collected from wild-type *R. delemar* hyphae in which further growth has been hampered by AmB concentrations ≥ 2 μg/ml showed little to no secretion/shedding of the toxin (**Extended Data Fig. 10**). These results confirm that the toxin is secreted/shed into the growth medium.

### The hyphae-associated toxin has structural features of ricin

Given the critical role of the toxin in the pathogenesis of mucormycosis, we did structural and bioinformatics studies to understand its mechanism of action. As reported in **Supplementary Table 1**, many of the toxin orthologues found in other organisms are annotated as ricin domain-containing proteins or ricin B chain-like lectins. Further detailed bioinformatic analysis of the *R. delemar* toxin sequence showed a two domain structure similar to that of ricin (Sequence ID: NP_001310630.1)^24^ (**Fig. 3a**). Specifically, the *R. delemar* toxin harbored a small region (amino acids 198-289) that resembled a sequence in ricin chain A, known to be involved in inactivating ribosomes (*i.e.* ribosome inactivating protein [RIP]), and two domains (amino acids 304-437 and 438-565) of the lectin-binding ricin chain B. Moreover, the *R. delemar* toxin contained an LDV-motif (**Fig. 3a**) that is present in ricin (**Fig 3a**, red colored amino acids) and that has been reported to cause damage to HUVECs *in vitro*^25,26^, and *in vivo* in models of vascular leak^27^ as well as postulated to cause ricin A chain-mediated vascular leak syndrome in humans^25^. Furthermore, gene ontology studies predicted that *R. delemar* toxin would have functions similar to ricin including sugar binding (GO:0005529, score of 0.64), as well as rRNA glycosylase activity (GO:0030598, score of 0.49) and hydrolase activity (GO:0004553, score of 0.35) (**Fig. 3a**, table).

**Figure 3.**
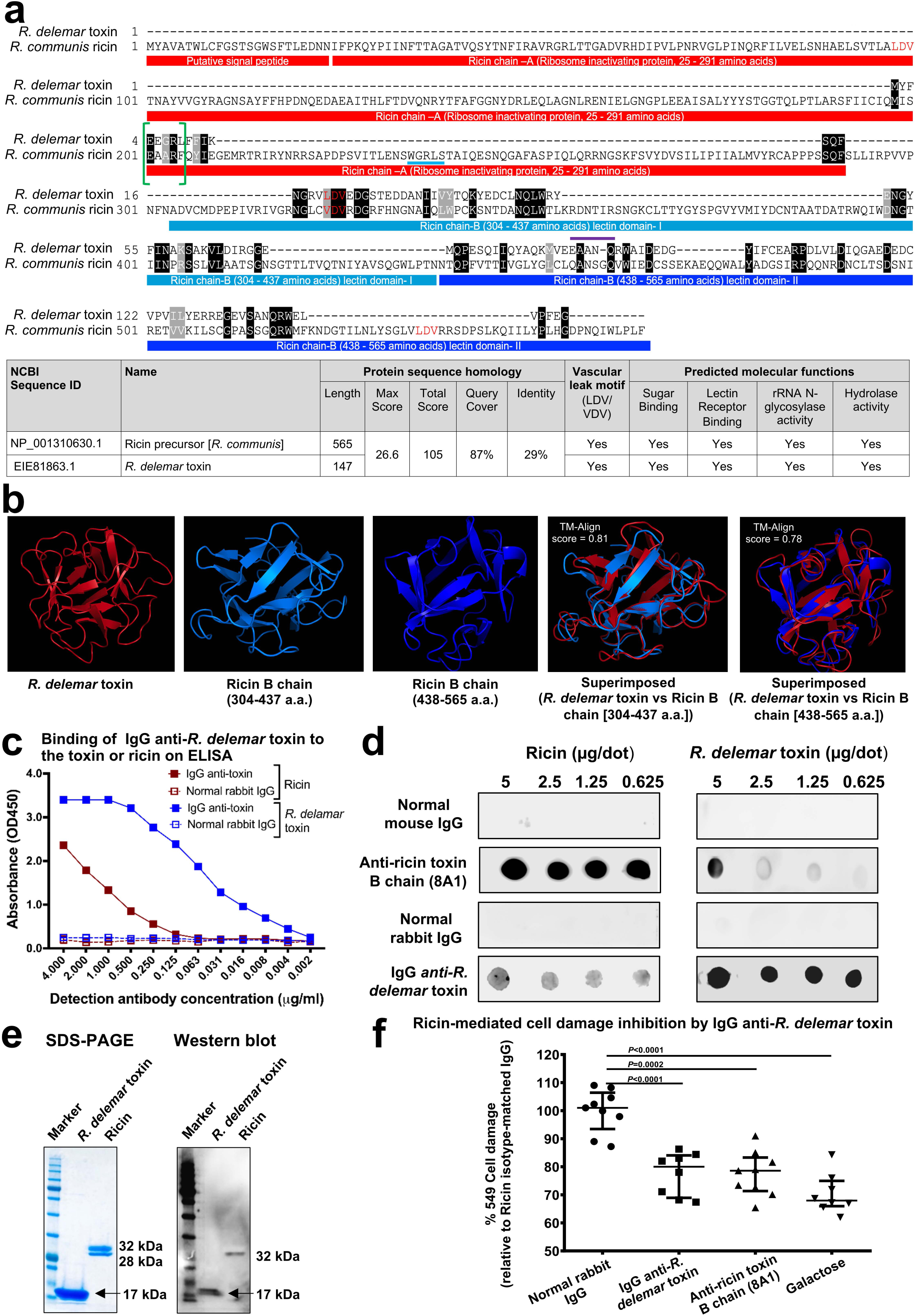
*R. delemar* toxin and ricin share structural features. **(a)** *R. delemar* toxin has 29% amino acid sequence identity with ricin. Both toxins share similar motifs and molecular functions. (**b)** 3-D structure model of *R. delemar* toxin shows similarities to ricin B chain. Protein 3D structure models of *R. delamar* toxin and ricin chain B (amino acids 304-437, and 338-565) were aligned residue-to-residue based on structural similarity using heuristic dynamic programming iterations and sequence independent TM-align score (0-1) were calculated based on structural similarity. TM-align score >0.5 considered significant similarity. **(c)** IgG anti-*R. delemar* toxin binds to ELISA plates coated with either *R. delemar* toxin or ricin. (**d**) Ricin is recognized on a dot blot by IgG anti-*R. delemar* toxin. (**e**) Western blot of *R. delemar* toxin and ricin using IgG anti-*R. delemar* toxin IgG. (**f)** IgG anti-*R. delemar* toxin, IgG anti-ricin (8A1 clone) (10 μg/ml each) or galactose (10 mM) inhibit ricin (77 nM)-mediated A549 cell damage (n=9 wells for normal rabbit IgG and Anti-ricin toxin B chain (8A1) group, n=8 for IgG anti-*R. delemar* toxin and galactose group pooled from three independent experiments). Data are median ± interquartile range. Statistical comparisons were made by using the non-parametric Mann-Whitney (two-tailed) test.

We produced a 3-D structural model of the *R. delemar* toxin by searching templates within the SWISS-Model template library (SMTL). The greatest resemblance was with sugar-binding proteins, especially galactose, the known lectin for ricin^28^. Other proteins with predicted resemblance to mucoricin included those with cell adhesion, toxin and hydrolase (glycosylase) activities (**Supplementary Table 2**). The rRNA glycosylase activity is a feature of ricin and is required for inactivating ribosomes^29^. Finally, the structures of the *R. delemar* toxin and ricin B chain were superimposable with a highly significant Tm-align score of 0.81and a score of 0.78 for ricin B domain I (amino acids 304-437) and domain II (438-565 amino acid), respectively (**Fig. 3b**). However, the 17 kDa *R. delemar* toxin is much smaller than either the A or B chains of ricin (32 kDa each). Nevertheless, the fungal toxin appears to share structural homology with portions of ricin that are responsible for inactivating ribosomes, inducing vascular leak and binding to galactose.

### *R. delemar* toxin is immunologically cross-reactive with ricin

To further explore the similarities between *R. delemar* toxin and ricin, we used the IgG anti-*R. delemar* toxin in an ELISA to determine whether the toxin and ricin were immunologically cross-reactive. Plates were coated with ricin or *R. delemar* toxin, and then incubated with the IgG anti *R. delemar* -toxin, or normal rabbit IgG. The former but not the latter bound to ricin or *R. delemar* toxin in a dose dependent manner (**Fig. 3c**). The IgG anti-*R. delemar* toxin also recognized ricin, and a murine monoclonal antibody (8A1) against the ricin B chain^30^ recognized the *R. delemar* toxin in a dot blot (**Fig. 3d**). Furthermore, the IgG anti-*R. delemar* toxin reacted with both the *R. delemar* toxin and ricin in a Western blot (**Fig. 3e**). Importantly, the IgG anti-*R. delemar* toxin protected A549 alveolar epithelial cells from ricin-induced damage in a manner similar to that of the IgG anti-ricin B chain (8A1 clone) or galactose [the lectin for the ricin chain B (**Fig. 3f**)]. Collectively, these data demonstrate the similarities between the two toxins.

### Mucoricin is a RIP that also promotes vascular permeability and induces both necrosis and apoptosis of host cells

After ricin is internalized by cells *via* its lectin-binding B chain, it’s A chain exerts its toxic activity by irreversibly inactivating ribosomes *via* its N-glycosylase activity. This result in the inhibition of protein synthesis^31^. The enzyme activity cleaves the N-glycosidic bond between the adenine nucleobase in the α-sarcin-ricin loop and its ribose causing the release of adenine (depurination)^32^. To determine whether the *R. delemar* toxin had similar activity, we compared the ability of the two toxins to inhibit protein synthesis in a cell-free rabbit reticulocyte assay^33^. As reported previously^34^, ricin concentration that caused 50% inhibition in protein synthesis (IC_50_) was ∼ at 2.2 x 10^−11^ M (**Fig. 4a**). The recombinant toxin of *R. delemar* also inhibited protein synthesis, albeit with ∼ 800-fold weaker activity than ricin (*i.e.* an IC_50_ of 1.7 x 10^−8^ M) (**Fig. 4b**). To determine if the protein inhibition is due to depurination, we used HPLC to measure the amount of adenine released from template RNA extracted from A549 alveolar epithelial cells^32^. In contrast to RNA exposed to the buffer control, RNA incubated with mucoricin released detectable amounts of adenine (**Fig. 4c**). The depurination is due to rRNA N-glycosylase activity since in contrast to the negative control, ovalbumin (OVA) which did not cleave 28S rRNA, the *R. delemar* toxin cleaved the rRNA in rabbit reticulocyte lysates after adding aniline, albeit at a concentration of 10^4^ higher than ricin and after incubation with the ribosomes for a longer period of time (**Fig. 4d**). Thus, like ricin, the *R. delemar* toxin is a RIP.

**Figure 4.**
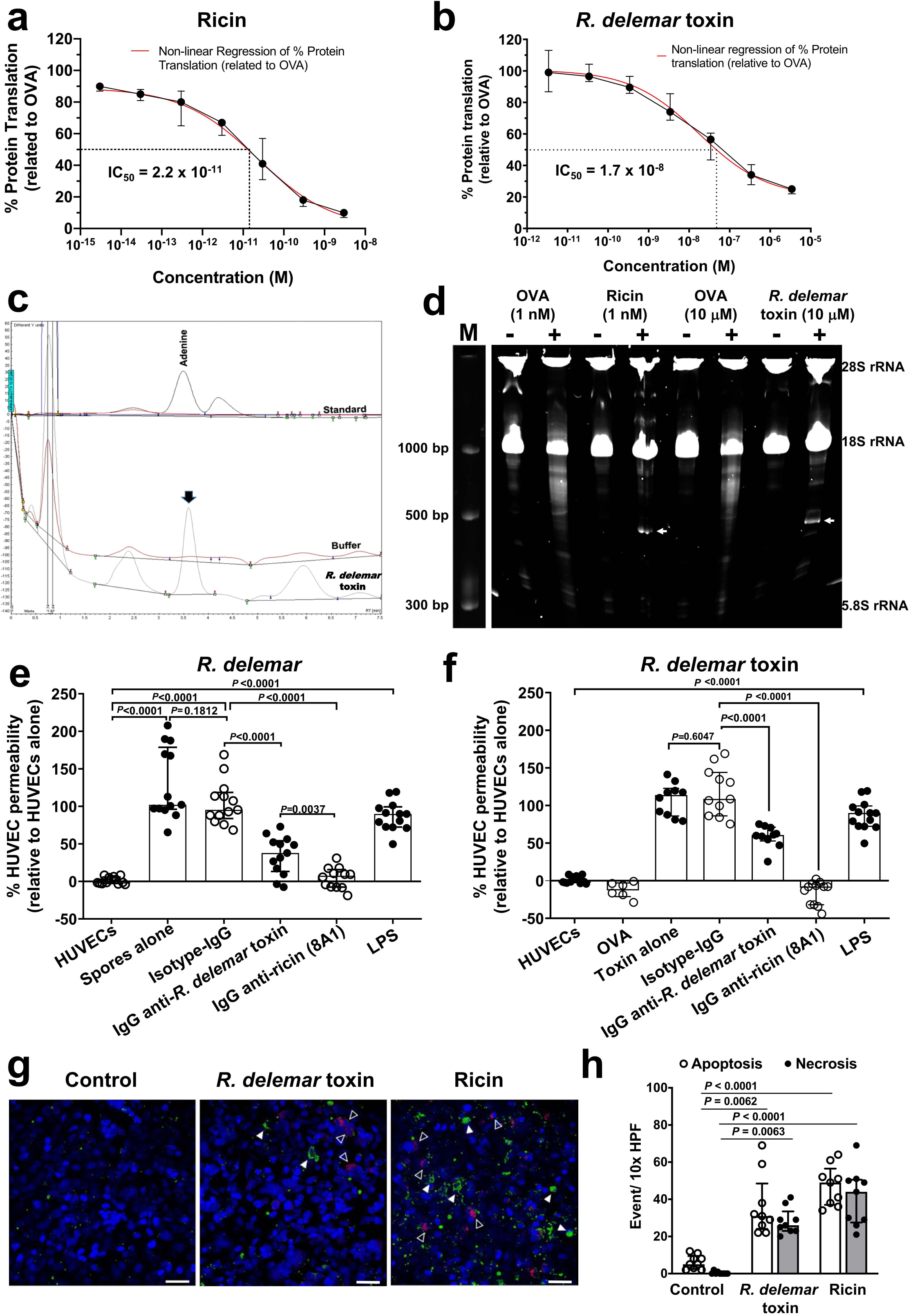
*R. delemar* toxin and ricin have functional similarities. **(a,b)** Cell-free rabbit reticulocyte assay showing protein synthesis inhibition by ricin (IC_50_ of 2.2 x 10^−11^ M) **(a)** and *R. delemar* toxin (IC_50_ of 1.7 x 10^−8^ M) **(b)**. Data (n=7 wells/concentrations for ricin; and n=6 wells/concentration for *R. delemar* toxin, pooled from three experiments) are presented as median ± interquartile range. **(c)** Representative HPLC chromatograms demonstrating the depurination activity of *R. delemar* toxin of A549 RNA at 3.6 min similar to adenine standard. (**d**) A representative gel (from three experiments) showing rRNA glycosidase activity of *R. delemar* toxin (1 μM) compared to ricin (1 nM) and control OVA (1 nM or 1 μM). Ribosomes were treated with ricin for 1 h or *R. delemar* toxin for 4 h. Extracted RNA were treated with (+) or without (-) aniline prior to running the gel. Arrows point to endo fragment at ∼500 bp. (**e, f**) *R. delemar* induces HUVEC permeability via its toxin. *R. delemar* (**e**) or recombinant toxin (2.9 μM) (**f**) were incubated with HUVEC for 5 h with or without 50 μg/ml of IgG isotype-matched or anti-*R. delemar* toxin or 10 μg/ml of IgG anti-ricin chain B (clone 8A1). LPS or OVA were added as a positive and negative controls, respectively. For **e,** n= 13 wells except for IgG anti-ricin 8A1 which n=12 wells pooled from three independent experiments. For **f**, n= 6 wells for Ova, n=10 wells for *R. delamar* toxin alone and *R. delamar* toxin + IgG anti-*R. delmar* toxin, n=11 wells for *R. delamar* toxin + Isotype IgG, n=12 wells for *R. delamar* toxin +IgG anti-ricin (8A1), and n=13 wells for HUVECs and LPS. Data in **e** and **f** were pooled from three independent experiments and presented as median ± interquartile range. (**g**) Detection of apoptosis/necrosis of A549 cells incubated for 2 h with 50 μg/ml (2.9 μM) of *R. delemar* toxin or 5 μg/ml (77 nM) ricin. Apoptotic cells (closed triangle) were identified by green fluorescence while necrotic cells (open triangle) are shown in red. Scale bar is 50 µm. (**h**) The number of apoptotic and necrotic events per high-power field (HPF) was determined, counting 10 HPF per coverslip. The data is combined from 3 independent experiments with each group in triplicate (total n=9 wells) and presented as median ± interquartile range. Kruskal-wallis test was used to compare control *vs. R. delamar* toxin or ricin.

In addition to its N-glycosylase activity, ricin chain A is known to cause vascular leak *in vitro*^25,26^ and *in vivo*^27^ mediated by its LDV sequence, a motif that is also present in the *R. delemar* toxin (**Fig. 3a**). To examine whether the *R. delemar* toxin compromised vascular integrity, we grew HUVECs on membrane inserts in transwells prior to treating the confluent monolayers with either *R. delemar* spores or *R. delemar* toxin for 5 h at concentrations that did not kill the HUVECs (**Supplementary Fig. 1).** The permeability of HUVECs was determined by measuring the amount of fluorescent dextran migrating from the upper chamber to the lower chamber of the transwells after adding *R. delemar* spores (**Fig. 4e**) or recombinant *R. delemar* toxin (**Fig. 4f**). Both *R. delemar* and the toxin induced permeability, which was equivalent to that induced by *E. coli* lipopolysaccharide (LPS), a potent inducer of vascular permeability^35^. Furthermore, IgG anti-*R. delemar* toxin blocked this enhanced permeability by 50-60%. Importantly, IgG anti-ricin B chain (8A1 clone) almost completely abrogated the permeability induced by *R. delemar* or its toxin (**Fig. 4e,f**). These results confirm that *R. delemar* induces permeability in HUVECs *via* its toxin and that IgG anti-ricin B chain blocks this *R. delemar*-mediated virulence trait.

Ricin is also known to cause cell damage by inducing both necrosis and apoptosis^36,37^. To determine whether the *R. delemar* toxin did the same, we used an Apoptosis/Necrosis detection kit to compare the abilities of *R. delemar* toxin and ricin to damage alveolar epithelial cells. As compared to the control, after 2 h both toxins caused comparable apoptosis and necrosis (**Fig. 4g,h**). Collectively, these results demonstrate the functional similarities between ricin and *R. delemar* toxin as RIPs that inhibits protein synthesis *via* N-glycosylase activity. Both toxins also cause cell death by apoptosis and necrosis. Based on the structural and functional similarity to ricin, we named the *R. delemar* toxin **mucoricin** and the corresponding gene **<u>R</u>icin-<u>L</u>ike <u>T</u>oxin (*RLT1***).

## Discussion

Mucormycosis is a lethal fungal infection often associated with extensive tissue damage. We have now identified a cell-associated/secreted/shed toxin that is widely present in pathogenic Mucorales fungi. We used genetic and biochemical techniques to show that the toxin gene, *RLT1* and its encoded protein (mucoricin) are required for the full virulence of *R. delemar.* In addition to being produced by pathogenic Mucorales, this toxin appears to be present in other fungi and bacteria. For example, orthologues in *Rhizophagus* were identified to have 30-40% identity with *RLT1*. *Rhizophagus* lives symbiotically with plants and is recognized as an integral part of the natural ecosystem and was shown to delay plant disease symptoms caused by *Phytophthora infestans*^38^. Similarly, *RLT1* orthologues were identified in the bacterial genera of *Streptomyces* and *Paenibacillus* (∼30% identity). Both bacteria are known inhabitants of soil, present in rhizosphere of various plants, and are used as biological control agents for crops because of their ability to secrete secondary metabolites^39^.

Mucoricin has structural and functional similarities to ricin, a prototypic Type 2 RIP consisting of two polypeptide chains (A and B) that are linked by a disulfide bond^24^. The A-chain of ricin is an *N-*glycosidase that is responsible for inactivating ribosomes, and the B chain is a galactose-specific lectin^29^ that enables the toxin to bind to target cells^28^. In contrast, Type I RIPs are monomeric A-chain-like RIPs^29^. Mucoricin appears to have the activities of both ricin A and B chains (and Type I RIPS) and both activities are present in a single 17 kDa protein (**Fig. 3e**). Several other RIPs isolated from plants consist of low molecular weight single chain proteins including a 26 kDa TRIP isolated from tobacco leaves^40^ and a 7.8 kDa protein isolated from sponge gourd seeds (*Luffa cylindrica*)^41^.

The strong sequence and structural similarity between mucoricin and ricin lies in the ricin B-chain, although mucoricin inhibits protein synthesis *via* N-glycosylase activity leading to depurination, albeit with lower activity than ricin. A possible explanation for the ability of mucoricin to inhibit protein synthesis is likely predicted by its conserved rRNA glycosidase activity. Specifically, the EAARF motif in ricin A chain is known to be responsible for the RIP activity of ricin. It is known to depurinate adenosine 4324 in 28S rRNA with the glutamic acid residue (E, shaded) responsible for this activity^42^. Furthermore, the arginine residue (R, shaded) separated by two amino acids from the glutamic acid residue (**Fig. 3a**, green brackets), is also required for the activity of the bacterial Shiga toxin, a potent ricin-like A chain with a fully conserved EAARF domain of ricin^43^. Mucoricin has the EEGRL, in which the glutamic acid and arginine residues are conserved (**Fig. 3a**, green brackets).

Another functional domain in ricin and Shiga toxin reported to be required for RIP activity is the WGRLS^44^ (**Fig. 3a**, cyan underline). This sequence also aligns with the EEGRL motif of mucoricin (**Fig. 3a**, green brackets; and **Supplementary Table 3**). Furthermore, mucoricin contains the EAANQ motif (**Fig. 3a**, purple overline) which resembles the ricin sequence of EAARF, with the glutamic acid residue conserved and the arginine residue replaced by asparagine, a conserved amino acid with properties that are weakly similar to arginine. The lack of fully conserved functional residues (*i.e*. arginine) between mucoricin and ricin/Shiga toxins likely explains the 800-fold weaker RIP activity of mucoricin as compared to ricin (**Fig. 4a,b**). Finally, mucoricin also has sequence homology with several other RIPs including saporin of *Saponaria officinalis,* a Type 1 RIP with 19% overall identity and 10 out of the 17 conserved amino acid residues in the functional domain of EAARF (**Supplementary Fig. 2**). Notably, the EEGRL and EAANQ are widely present in Mucorales fungi known to cause human disease (**Supplementary Table 3**). Thus, mucoricin appears to be a RIP that functionally resembles ricin A chain and Type-1 RIPs such as saporin^45^. The contribution of the EEGRL and EAANQ motifs to the RIP activity is being further investigated.

Our *in vitro* studies suggest that *RLT1* is expressed most strongly when the hyphal matt is aeriated and in response to alveolar epithelial cells (**Extended Data Fig. 4c,d**). These results further suggest that mucoricin may be highly active during pulmonary mucormycosis and potentially in rhinoorbital disease, when hyphae are exposed to epithelial cells in the presence of ambient levels of oxygen. It is of interest that, mitochondria are believed to play a central role in RIPs’ (e.g. ricin, Shiga toxin and abrin) ability to induce host cell apoptosis^46^. Although the expression of *RLT1* in *R. delemar* is higher in response to alveolar epithelial cells than to HUVECs (**Extended Data Fig. 4d**), the latter host cells are damaged much more rapidly by *R. delemar* (*i.e.* significant *R. delemar-*induced HUVEC injury occurs at 8 h *vs.* 48 h for *R. delemar-*induced alveolar epithelial cells [**Extended Data Fig. 1**])^13^, and by purified mucoricin (**Fig. 1a**). In accord with these results, it has been shown that HUVECs are rapidly damaged and their permeability affected by small peptides containing the LDV-motif but lacking the sequences responsible for N-glycosidase activity^26^. In this study we show that *R. delemar* compromises the permeability of HUVEC monolayers *in vitro* by the direct effect of mucoricin. The LDV and other (x)D(y) motifs (with known vascular leak effector function) are widely present in pathogenic Mucorales **(Supplementary Table 3**)^25^. Thus, it is possible that the LDV-motif is responsible for angioinvasion and rapid hematogenous dissemination in mucormycosis^6^ by inducing damage to vascular endothelial cells.

The exact mechanism by which mucoricin enters a target cell to exert its lethal effect is not yet known. However, our data strongly indicate that it is cell-associated as well as secreted/shed by Mucorales. The amount of toxin in the medium (27 nM, **Extended Data Fig. 10b**) from a small-scale growth in a 96-well plate was sufficient to exceed the IC_50_ in RIP activity of 17 nM (**Fig. 4b**). Thus, secreted/shed toxin likely exerts its toxicity by binding to and then entering the host cells in the absence of invading hyphae. Alternatively, invading *R. delemar* hyphae release the toxin once they are phagocytosed by immune^9^ cells or barrier cells^13,47,48^.

Our *in vivo* studies clearly demonstrate the contribution of mucoricin to pathogenesis by enhancing angioinvasion, inflammation and tissue destruction. There is also evidence that the lethality of ricin *in vivo* is related at least in part to its ability to induce a massive inflammatory immune response accompanied by infiltration of PMN^49^ in many settings such as acute lung injury^49,50^ and gastrointestinal disease^51^. This is likely due to activation of the innate arm of the immune system by the toxin itself or by toxin-damaged cells. Indeed, our histopathological examination of organs harvested from mice injected with purified mucoricin shows inflammation and recruitment of PMNs (**Fig. 1e**). Neutralizing the effect of the toxin either by RNAi or by anti-mucoricin antibodies decreased inflammation and host tissue damage (**Fig. 2e-h** and **Extended Data Fig. 8**). These results confirm the critical role of mucoricin in the pathogenesis of mucormycosis and suggest that it is involved in mediating inflammation and tissue damage, both of which are clinical features of mucormycosis. Of note, treatment of mucormycosis patients with antifungal agents is often hampered by the extensive tissue necrosis that prevents optimal delivery of drugs into the site of infection. Hence, antifungal treatment alone (without surgical intervention) is often non-curative^6^. Thus, antibody-mediated neutralization of mucoricin might reduce tissue necrosis, decrease the need for disfiguring surgery, and maximize the effect of antifungal therapy.

Based on these results, we propose a model of pathogenesis and the role of mucoricin in this process. We suggest the following events.

i. Infection is initiated when fungal spores are inhaled, and in the absence of phagocytes (or presence of dysfunctional phagocytes such as in patients with DKA). Fungal spores express CotH^10–12^ and bind to either GRP78 on nasal epithelial cells^48^, or to integrin β1^48^ which activates epidermal growth factor receptor (EGFR)^47^ on alveolar epithelial cells to induce invasion.
ii. Under aerobic conditions, the calcineurin pathway is activated in the inhaled spores, causing them to germinate^52,53^, a process that leads to the production of mucoricin.
iii. Mucoricin binds to tissue cells by its lectin receptor, inhibits protein synthesis, and causes apoptosis and necrosis. The toxin can also compromise vascular permeability resulting in rapid hematogenous dissemination and tissue edema often seen in patients with mucormycosis.
iv. While tissue damage is occurring, and because the toxin and debris from necrotic cells are recognized by the immune system, an inflammatory immune response leads to the recruitment of PMNs and other tissue-resident phagocytes.
v. Although the recruited phagocytes damage some of the invading hyphae, both the dead and live hyphae release mucoricin, resulting in more host cell death and more inflammation.
vi. In the necrotic tissue, the fungus can proliferate, protected from both phagocytes and anti-fungal drugs.

Our finding that mucoricin remains active even in dead organisms offers an explanation for why antifungal therapy alone has limited efficacy in patients with mucormycosis, and why the fungal lesions must frequently be surgically excised. Importantly, other toxins/mechanisms of host cell damage likely exist in Mucorales, since antibody blocking studies and downregulation of toxin gene expression do not <u>fully</u> abrogate the ability of *R. delemar* to damage host tissues.

In summary, we have identified a ricin-like toxin (mucoricin) that is widely present in Mucorales fungi, where it plays a central role in the pathogenesis of mucormycosis. We postulate that strategies to neutralize mucoricin will have significant therapeutic benefits.

Correspondence and requests for materials should be addressed to Ashraf S. Ibrahim

## Acknowledgments

This work was supported by a Public Health Service grant R01AI063503 and R01AI141360 to ASI. MS is supported by R00DE026856, ESV by R01A11752861, VMB by U19AI110820 and R01AI141360, and SGF by R01AI124566 and R01DE022600. ESV is also supported by the Simmons Patigian Distinguished Chair and a Distinguished Teaching Chair. AR is sponsored by the SURF program at UT Southwestern and is currently at Vanderbilt University.

We would like to thank Dr. Samuel French for his assistance in reading the histopathological slides of the purified mucoricin, Ms. Heewon Jeon, Ms. Ayesha Ahmed, and Mr. Stephen Ruback for technical assistance. We thank Dr. David Vance and Ms. Greta Van Slyke for their work on the 8A1 monoclonal antibody and Dr. Robert Munford, NIH for his insightful suggestions concerning the nature of the toxin.

## Author contributions

S.S.M.S. conceived, designed and performed studies to purify and identify the toxin, and screen its activity *in vitro* and *in vivo* and wrote the manuscript. C.B. generated mucoricin mutants and characterized their virulence *in vitro* and *in vivo* and conducted the antibody efficacy studies. Y.G. helped in animal studies, conducted confocal microscopy, cross reactivity studies, and RIP activity studies. S.S. designed and performed homology modeling, cross reactivity studies, and toxin secretion studies. T.G. helped in the animal studies. M.S. performed the necrosis/apoptosis assay and the mouse immunohistochemistry studies. A.A. performed permeability studies, E.G.Y. performed sequence alignment and gene ontology studies. S.A. purified recombinant toxin and polyclonal antibodies. A.P. and G.C. provided and performed the human immunohistochemistry studies. C.P. and V.V. performed and interpreted the mouse histology studies. AR carried out studies on cross-reactivity of mucoricin and ricin. V.M.B. and J.D.H. performed phylogenetic studies and blast search of mucoricin in Mucorales. N.J.M. generated and characterized the 8A1 monoclonal antibody. J.E.E. and S.G.F. provided intellectual advice, designed studies, and edited the manuscript. E.S.V. conceived, designed and carried out studies of cross reactivity, provided reagents and expertise on ricin and helped write the manuscript. A.S.I. conceived, designed, coordinated and supervised the studies, performed experiments, analyzed data, and wrote the manuscript along with comments from co-authors.

## Materials and Correspondence

Reprints and permissions information is available at www.nature.com/reprints

## Competing interests

A.S.I. owns shares in Vitalex Biosciences, a start-up company that is developing immunotherapies and diagnostics for mucormycosis. The remaining authors declare no competing interests.

The Lundquist Institute has filed intellectual property rights concerning mucoricin. Vitalex Biosciences has an option to license the technology from The Lundquist Institute for Biomedical Innovation.

Other authors declare no conflict of interest.

Correspondence and requests for materials should be addressed to.

## Methods

### Organisms, culture conditions and reagents

*R. delemar* 99-880 and *R. oryzae* 99-892 were isolated from the brain and lungs of patients with mucomycosis and obtained from the Fungus Testing Laboratory, University of Texas Health Science Center at San Antonio which had its genome sequenced^19^. *Cunninghamella bertholletiae* 182 is a clinical isolate and is a kind gift from Thomas Walsh (Cornell University). *Lichtheimia corymbifera* is also a clinical isolate obtained from the DEFEAT Mucor clinical study^54^. *R. delemar* M16 is a *pyrf* null mutant derived from *R. delemar* 99-880 and was used for transformation to attenuate mucoricin expression^55^. The organisms were grown on potato dextrose agar (PDA, Becton Dickinson) plates for 5-7 days at 37°C. For *R. delemar* M16, PDA was supplemented with 100 mg/ml uracil. The sporangiospores were collected in endotoxin free Dulbecco’s phosphate buffered saline (PBS) containing 0.01% Tween 80, washed with PBS, and counted with a hemocytometer to prepare the final inoculum. To form germlings, spores were incubated in YPD (Becton Dickinson) medium at 37°C with shaking for different time periods. Finally, for growth studies 10^5^ spores of *R. delemar* wild-type, or mutant strains were plated in the middle of PDA agar plates. The plates were incubated at 37°C and the diameter of the colony was calculated every day for 6 days. The monoclonal anti-ricin B chain antibody (clone 8A1)^30^ and affinity purified rabbit anti-ricin antibodies were prepared and characterized in the Vitetta and Mantis laboratories. Galactose was obtained from Fisher Scientific (Cat # BP656500) and used for blocking the damaging effect of ricin holotoxin.

### Host cells

Human alveolar epithelial cells (A549 cells) were obtained from a 58-year-old male Caucasian patient with carcinoma and procured from American Type Culture Collection (ATCC). The cells were propagated in F-12K Medium developed for lung A549 epithelial cells. Primary alveolar epithelial cells were obtained from ScienCell (Cat # 3200) and propagated in Alveolar Epithelial Cell Medium (Cat#3201) and passaged once.

HUVECs were collected by the method of Jaffe *et al.*^56^. The cells were harvested by using collagenase and were grown in M-199 (Gibco BRL) enriched with 10% fetal bovine serum, 10% defined bovine calf serum, L-glutamine, penicillin, and streptomycin (all from Gemini Bio-Products, CA). Second-passage cells were grown to confluency in 24- or 96-well tissue culture plates (Costar, Van Nuys, CA) on fibronectin (BD Biosciences). All incubations were in 5% CO_2_ at 37°C. The reagents were tested for endotoxin using a chromogenic limulus amebocyte lysate assay (BioWhittaker, Inc.), and the endotoxin concentrations were less than 0.01 IU/ml.

Fresh red blood cells were isolated from blood samples collected from healthy volunteers after obtaining a signed informed consent form and processed as previously described^57^. Endothelial cell and red blood cell collection was approved by Institutional Review Board at The Lundquist Institute at Harbor-UCLA Medical Center.

### Purification and characterization of ricin

Two sources of ricin were used. One was purified from a large stock of pulverized castor beans in the Vitetta laboratory^58^. Its IC_50_ and toxicity were tested on Daudi lymphoma cells (ATCC® CCL-213™), HUVECs, and in cell free reticulocyte assays^25,34,59^. The other source was purchased from Vector Laboratories (Burlingame, CA; Cat No. L-1090). Both sources were similar in purity and activity.

### Purification and identification of mucoricin

To purify mucoricin, *Rhizopus* fungal spores (10^3^/ml) were grown for 5 days at 37°C in YPD culture medium. The supernatant was separated from the fungal mat by filtration and the fungal mycelia was ground in liquid nitrogen and extracted with sterile water, concentrated and analyzed through size exclusion columns^15^. Host cell damage assay showed that substances >10 kDa and < 30kDa is responsible for injuring the host cells (**Extended Data Fig. 1a**). The concentrated water extract was then subjected to 3D chromatographic separations (**Extended Data Fig. 1**). For the first dimension, the concentrated extract run on native polyacrylamide gel under electrophoretic force (**Extended Data Fig. 1b**). The gel was cut into 6 pieces and then eluted separately in PBS buffer prior to testing for their damaging activity. Only fraction #6 corresponding to 15-20 kDa showed damaging effect to host cells (**Extended Data Fig. 1c**). Next, this fraction was concentrated and subjected to separation using methanol:water (4:1) as a solvent on cellulose plates. Fractions were scraped from the cellulose plate and dissolved in irrigation water, followed by incubation with host cell after filter sterilization. A third dimensional fractionation was applied to the fraction that caused damage using cellulose plates and with the solvent system as above. This round of fractionation resulted in one fraction causing damage to host cells.

### Structural modeling of mucoricin

For amino acid sequence comparisons, mucoricin and ricin (Sequence ID: NP_001310630.1) protein sequences were aligned using MUSCLE/CLUSTAL-W. We searched the SWISS-MODEL template library (SMTL) (https://swissmodel.expasy.org/) to find templates for building 3-D structural model of mucoricin. Briefly, a BLAST search of the SMTL against the primary amino acid sequence identified the target sequence. To build the model, we performed target-template alignment using ProMod3, and templates with the highest quality were selected for model building. Insertions and deletions were re-modeled using a fragment library, and the side chains were rebuilt. Finally, the geometry of the resulting model is regularized by using a force field. In case loop modeling with ProMod3 fails, an alternative model is built with PROMOD-II. The models showing high accuracy values were finalized for similarity comparisons. Ricin 3-D structure models were also built using ricin chain B amino acids 313-435 and 440-565^60–62^. Ricin and mucoricin 3-D protein models were aligned using MacPyMOL software and Tm align score was calculated by webtool Tm Align (https://zhanglab.ccmb.med.umich.edu/TM-align)63. Using gene ontology term prediction, mucoricin was predicted to have carbohydrate-binding, hydrolase activity and negative regulation of translation functions.

### Expression and purification of mucoricin

Heterologous expression of mucoricin gene in *S. cerevisiae* was performed to ensure the production of a functional toxin since we used this yeast to generate functional *R. delemar* proteins before^11,64^. The heterologous expression was conducted as follows; total RNA was isolated from *R. delemar* hyphae grown on YPD broth and reverse transcribed into cDNA. The entire ORF of mucoricin was PCR amplified from cDNA using Phusion High-Fidelity PCR Kit (New England Biolabs) using the primers 5’-GATAAG<u>ACTAGT</u>ATGTATTTCGAAGAAGGC-3’ and 5’-GGTGATG<u>CACGTG</u>TCCTTCAAATGGCACTA-3’. The amplified PCR product was verified by sequencing and then cloned into modified XW55 yeast dual expression vector^65^ in the highlighted *Spe*I and *Pml*I sites downstream of the ADH2 promoter by yeast recombinase technology [protocolYeastmaker™ YeastTransformation System 2 (Clontech)] and according to the manufacturer’s instructions. The generated yeast expression vector was transformed into *S. cerevisiae* strain BJ5464 using protocol Yeastmaker™ Yeast Transformation System 2 (Clontech). The transformants were screened on yeast nitrogen base (YNB) medium lacking uracil. *S. cerevisiae* transformed with empty plasmid was served as negative control. Transformants were grown on YNB without uracil for 1-3 days then transformed into YPD medium for 3 days at 30°C with shaking. The expression of mucoricin was induced once the glucose was exhausted from the medium and yielded a recombinant mucoricin that was both 6x His- and Flag-tagged. Purification of the recombinant mucoricin was performed by Ni-NTA matrix affinity purification according to the manufacturers’ instructions (Sigma-Aldrich). The purity of the protein was confirmed by SDS-PAGE and quantified by a modified Lowry protein assay (Pierce).

### Anti-mucoricin and anti-ricin antibodies

Rabbit and mouse monoclonal antibodies against recombinant mucoricin coupled to KLH were raised by ProMab Biotechnologies Inc.^11^. The IgG fraction was purified from the antisera using protein A/G spin column (Thermo Fisher Scientific) according to the manufacturer’s instructions. Normal IgG was purified from the sera of non-immunized rabbits and used as a control. Rabbit and monoclonal antibodies against ricin and ricin B chain were prepared as described previously^27,30^. Hybridoma cells producing anti-muroricin antibodies were propagated in WHEATON CELLine bioreactor 350 using protein-free hybridoma medium 1× (Gibco) for 5 to 7 days at 37°C in 5% CO_2_. The antibody-containing supernatant was collected and purified using protein G spin columns (Thermo Fisher Scientific). Eluted purified IgG were dialyzed in PBS using a dialysis cassette (Thermo Fisher Scientific), and the purity of the antibodies was confirmed by SDS-PAGE prior to determining the concentration using the Bradford protein assay (Bio rad, Hercules, CA). Endotoxin levels were measured by the Limulus Amebocyte Lysate (LAL) kit (Charles River) and determined to be <0.8 EU/ml which is below the 5 EU/kg body weight set for intraperitoneal injection^66^. Mouse monoclonal antibody clone 8A1 was raised against the ricin B chain^30^, which has 33% sequence identify with mucoricin. Clone 8A1 recognizes an epitope mapped to ricin B chain (2γ, amino acids 221-262).

### Cell damage assay

The damage of epithelial cell [A549 and Primary (ScienCell, Cat # 3200)] and HUVECs was quantified using a ^51^Cr-release assay^67^. Briefly, confluent cells grown in 24-well tissue culture plates were incubated with 1 μCi/well Na_2_^51^CrO_4_ (ICN) in F12K-medium (for epithelial cells) or M-199 medium (HUVECs) for 16 h. On the day of the experiment, the media was aspirated, and cells were washed twice with pre-warmed Hanks’ balanced salt solution (HBSS, ScienCell). Cells were treated with toxin suspended in either 1 ml of F12K-medium (for epithelial cells) or RPMI 1640 medium (for endothelial cells) supplemented with glutamine and incubated at 37°C in a 5% CO_2_ incubator. Spontaneous ^51^Cr release was determined by incubating the untreated cells in the same volume of the culture medium supplemented with glutamine. At different time points, and after data were corrected for variations in the amount of tracer incorporated in each well, the percentage of specific cell release of ^51^Cr was calculated as follows: [(experimental release) – (spontaneous release)]/[1 – (spontaneous release)]^68^. Each experimental condition was tested at least in triplicate, and the experiment was repeated at least once.

In some experiments, the effect of mucoricin gene silencing on damage to HUVECs or A549 cells was measured by incubating 1.0 x 10^6^/ml or 2.5 x 10^5^/ml spores of *R. delemar* and incubated for 6 or 48 h, respectively. In other experiments the protective effect of IgG anti-mucoricin was measured by incubating the fungal cells with either 50 μg/ml of IgG anti-mucoricin or normal rabbit IgG (R & D Systems, Cat # AB-105-C) for 1 h on ice prior to adding the mixture to A549 cells radiolabeled with ^51^Cr. The assay was carried out for 48 h. The amount of damage was quantified as above.

To study the effect of fungal cell viability on host cell damage, fungal spores (10^6^/ml) were cultured in F12K media and left to grow overnight at 37°C. The fungal hyphae were collected by filtration, dried by padding with a sterile filter paper, weighed and then aseptically cut into four equal small pieces of 0.1 mg wet weight. The fungal hyphal matt was suspended in 1 ml F12K and heated at 60°C in a water bath for 4 h and then cooled down. To check the viability of the hyphal matt, a loop full of the hyphae was plated on PDA plates. The other two groups of fungal hyphae were suspended in preheated and cooled in F12K culture media. Another group of F12K culture media was prepared by heating at 60°C and then cooled to represent spontaneous control. The fungal samples were incubated with ^51^Cr-labelled A549 alveolar epithelial cells previously seeded into 24-well plates as above and the damage assay was carried out for 24 h at 37°C and the amount of ^51^Cr released in the supernatant was measured as above.

To determine whether IgG anti-mucoricin protected cells against ricin-induced damage, 5 μg/ml (∼77 nM) of ricin was incubated with either 10 μg/ml of the monoclonal IgG anti-ricin B chain (clone 8A1)^69^, 10 μg/ml of IgG anti-mucoricin, or normal rabbit IgG (R & D Systems, Cat # AB-105-C) or 10 mM of galactose on ice for 1 h prior to adding to ^51^Cr-labelled confluent A549 alveolar epithelial cells in 24-well plate. The damage assay was conducted as above for 24 h.

### Western blotting

Hyphal expression of mucoricin was determined in *R. delemar* wild-type, or RNAi mutants from hyphal matt grown for overnight at 37°C in YNB without uracil medium^70^. Briefly, mycelia were collected by filtration, washed briefly with PBS, and then ground thoroughly in liquid nitrogen using mortar and pestle for 3 min. The ground powder was immediately transferred to microfuge tube containing 500 µl extraction buffer which consisted of 50 mM Tris-HCl, pH 7.5, 150 mM NaCl, 10 mM MgCl_2_. The extraction buffer was supplemented with 1X Halt Protease Inhibitor Cocktails (Thermo Scientific) and 1 mM phenylmethylsulfonyl fluoride (PMSF). The sample was vortexed vigorously for 1 min, then centrifuged for 5 min at 21000 g at 4°C. The supernatant was filtered with PES syringe filters (Bioland Scientific, Cat# SF01-02) and transferred to a new tube and the protein concentration determined using Bradford method.

For Western blotting, 10 µg of each sample was used to separate proteins on an SDS-PAGE. Separated proteins were transferred to PVDF membranes (GE Water & Process Technologies) and treated with Western blocking reagent (Roche) for overnight at 4°C. The IgG anti-mucoricin (2 μg/ml) and the murine 8A1 monoclonal anti-ricin B chain were used as primary antibodies. After 1 h, 0.5 μg/ml of HRP-IgG anti-rabbit IgG (Jackson ImmunoResearch, Product number 111-035-144) (when rabbit IgG anti-mucoricin was used as a primary) or HRP-IgG anti-mouse (Invitrogen, Cat #31450) (when murine 8A1 antibody anti-ricin B chain used as a primary) secondary antibodies (Jackson Immuno Research) were added for another 1 h at room temperature. Mucoricin bands were visualized by adding the HRP substrate (SuperSignal West Dura Extended Duration Substrate, Thermo Scientific), and the chemiluminescent signal was detected using an In-gel Azure Imager c400 fluorescence system (Azure Biosystems). The intensity of the bands was quantified by ImageJ software.

To examine the cross reactivity of ricin with IgG anti-mucoricin, 5 µg of ricin (77 pmol) or recombinant mucoricin (294 pmol) were submitted to SDS-PAGE under reducing and denaturing conditions. Western blotting was conducted as above using IgG anti-mucoricin as a primary followed by HRP-IgG anti-rabbit IgG as secondary (Jackson ImmunoResearch, Product number 111-035-144).

### Gene expression analysis

Expression of the mucoricin gene was studied in *R. delemar* as spores (10^3^/ml) germinated into hyphae in YPD medium for 16 h at 37°C. At selected times, cells or mycelia were collected by centrifugation, followed by filtration using 0.22 μm membrane units. The cells were washed once with PBS, and ground in liquid nitrogen using mortar and pestle. RNA was extracted using RNeasy Plant Mini kit (Qiagen). To quantify the expression of the mucoricin gene in response to host cells, fungal spores (10^6^/ml) were incubated with either epithelial, HUVECs or human erythrocytes in 24-well plate using F12K, RPMI, or PBS, respectively. The fungal cells were collected at different time intervals including zero, 2 and 5 h and directly ground with liquid nitrogen followed by RNA extraction using RNeasy Plant Mini Kit. Contaminating genomic DNA was removed from RNA samples by treatment with 1 μl of Turbo-DNase (Ambion) for 30 min at room temperature. DNase was then removed using an RNA Clean-Up kit (Zymo Research). First-strand cDNA synthesis was performed using the Retroscript first-strand synthesis kit (Ambion). Toxin specific primers were designed with the assistance of online primer design software (Genscript). Mucoricin gene primers include G7F1; 5’-CTGGCGTTACGAAAATGGTT-3’ and G7R1; 5’-TAAATCAGGACGGGCTTCAC-3’. The amplification efficiency was determined by serial dilution experiments, and the resulting efficiency coefficient was used for the quantification of the products ^71^. Gene expression was analyzed by an ABI Prism 7000 Sequence Detection System (Applied Biosystems) using the QuantiTect Sybr Green PCR kit (Qiagen). PCR conditions were 10 min at 90°C and 40 cycles of 15 s at 95°C and 1 min at 60°C. Single PCR products were confirmed with the heat dissociation protocol at the end of the PCR cycles. The amount of gene expression was normalized to actin [ACT1-RT5′; 5’-TGAACAAGAAATGCAAACTGC-3’ and ACT1-RT3′; 5’-CAGTAATGACTTGACCATCAGGA-3’] and then quantified using the 2(−ΔΔC(T)) method ^72^. All reactions were performed in triplicate, and the mixture included a negative no-reverse transcription (RT) control in which reverse transcriptase was omitted.

### *In vitro* apoptosis/necrosis assay

A549 lung epithelial cells were grown to confluency on fibronectin-coated circular glass coverslips in 24-well tissue culture plates and then incubated with 50 μg/ml (2.9 μM) mucoricin or 5 μg/ml (77 nM) ricin (concentrations shown to cause *in vitro* damage to alveolar epithelial cells [**Fig. 3f**]) for 2 hours after which the cells were washed and stained with 1x Apoxin Green Indicator and 1x 7-AAD (Apoptosis/Necrosis detection kit, Abcam) for 45 min. The cells were fixed and mounted in ProLong Gold antifade containing DAPI (Life Technologies) to visualize cells. Microscopic z-stack pictures were taken using a Leica SP8confocal laser scanning platform. Apoptotic cells *vs.* necrotic cells were identified by their green and red fluorescence, respectively. The number of apoptotic and necrotic events per high-power field (HPF) was determined, counting 10 HPF per coverslip. The experiment was performed three times in triplicate.

### *In vitro* protein translation assay

The ability of the two toxins to inhibit protein synthesis was measured by using a modification of a previously described method^33^. Briefly, a rabbit reticulocyte lysate (Promega, Cat: L4151) was thawed at 37°C immediately before use and supplemented with 40 μl of 1 mM hemin stock solution and 10 μl of 1 M creatine phosphate (Sigma-Aldrich, Cat: 27920) and 10 μl of 5 mg/ml creatine phosphokinase (Sigma-Aldrich, Cat:C7886) before the lysate had fully thawed. The reaction mixture was prepared into 96-well plates as follows: 1 μl of 1 mM amino acid mixture minus methionine (Promega, Cat: L9961), 35 μl of rabbit reticulocyte lysate, 1 μl of 7-fold dilutions of ricin, mucoricin, control OVA, or cycloheximide (Fisher Scientific, Cat: AC357420010). Diluted distilled water was added to a final volume of 48 μl. Two replicates were employed in all experiments and the experiments were repeated at least three times. After a pre-incubation period of 30 min at 37°C, 2 μl ^35^S Methionine (1,200 Ci/mmol) (PerkinElmer) was added to a final volume of 50 μl. The 96-well plate was incubated at 30°C for 60 min. Two μl from each well was added per well of a 24-well plate containing 98 μl of 0.5 M H_2_O_2_. Proteins were precipitated with 900 μl of 25% trichloroacetic acid (TCA) before harvesting precipitates on Whatman filter strips (Sigma-Aldrich, Cat: WHA1823035). Filter paper disks were placed in Biofluor scintillation fluid (Perkin Elmer, Cat: 6013329), and [^35^S] Methionine incorporation was quantitated by scintillation counting. Background counts determined from well containing all reagents without rabbit reticulocyte lysate were subtracted from all CPMs.

### The depurination activity assay of mucoricin

The depurination activity of mucoricin was measured by the release of adenine when mammalian RNA was treated with mucoricin for 24 h at 37°C^32^. Mammalian RNA extracted from A549 alveolar epithelial cells by Qiagen RNasy mini kits according to manufacturer’s instruction were treated with 20 µg/ml of mucoricin in 0.01M HEPES/10 mM ammonium acetate buffer containing 1 mg/ml BSA for 24 h at 37°C. The solution was then filtered through a 10 kDa size exclusion column and 40 µl was injected into HPLC using Phenomenex Luna C18 reverse phase column (10 x 250 mm) attached to a Varian ProStar HPLC 218 system (Varian, Walnut Creek, CA). Solvent A was 20 mM ammonium acetate, and solvent B was 100% acetonitrile. The column gradient was as follows: 97% to 60% solvent A in 10 min at flow rate of 1 ml/min. The column effluent was monitored at 260 nm.

### Glycosylase activity assay

The N-glycosylase activity of toxins were determined by using rabbit reticulocyte lysate^73,74^. Briefly, 40 μl of lysate was incubated with ricin (1 nM), mucoricin (10 μM) or control OVA (1 nM or 10 μM) in the presence of 10 mM MgCl_2_ at 30°C for 1 or 4 hours. After the treatment, ribosomal protein was denatured by 50 mM Tris/0.5% SDS to release RNA. The RNA was purified by phenol-Tris extraction followed by ethanol precipitation. Half of each purified RNA sample was subjected to 2 M aniline acetate (pH 4.5) treatment for 10 minutes on ice, while the other half of RNA was incubated without aniline treatment. rRNA were further extracted using water saturated ether followed by ethanol precipitation. Three micrograms of each rRNA sample were resolved on 7 M Urea Polyacrylamide gel electrophoresis and RNA fragment bands were visualized by staining with ethidium bromide.

### Transwell permeability assay^75^

HUVECs were seeded on 24-Corning transwell plate with permeable polyester inserts (0.4 μm, Fisher) coated with fibronectin (15 µg/ml in PBS, Fisher). HUVECs were grown to confluency in M-199 medium with phenol red. *R. delemar* spores (10^5^) in M-199 (without phenol red) were added to the upper chamber, and the plate was incubated for 5 h at 37°C. As a positive control for the permeability of HUVECs, *E. coli* LPS (Sigma-Aldrich) was added at 2 μg/ml to uninfected HUVECs. Following incubation, 3 µl of 50 mg/ml FITC-dextran-10K (Sigma) was added to the upper chamber of the trans-well and the migration of the dextran through the HUVEC monolayer to the lower trans-well was determined 1 h later by quantifying the concentration of the dye in the bottom chamber using florescence microplate reader at 490 nm^75^. To determine the direct effect of mucoricin on the permeability of the HUVEC monolayer, 50 μg/ml (2.9 μM) mucoricin or control OVA were added to the HUVEC-seeded wells instead of *R. delemar*. To determine the effect of antibodies on permeability induced by *R. delemar* or mucoricin, 50 μg/ml of normal rabbit IgG (R & D Systems, Cat # AB-105-C), IgG anti-mucoricin, or 10 μg/ml of IgG anti-ricin toxin chain B (clone 8A1) were incubated for 30 min on ice with *R. delemar* spores or mucoricin prior to their addition to the upper chamber of the trans-well.

### *In vivo* effects induced by mucoricin

To test the effect of the purified toxin *in vivo*, male (ICR mice, ∼27-32g) were immunosuppressed by intraperitoneal injection of 200 mg/kg of cyclophosphamide and subcutaneous injection of 250 mg/kg cortisone acetate on day −2 and +3, relative to toxin injection. This regimen results in approximately 10 days of leucopenia with reduction in neutrophils, lymphocytes and monocytes as described previously^76^. Mouse gender has no effect on the pathogenesis of mucormycosis, or antifungal treatment as determined by an NIH Contract No. HHSN272201000038I/Task Order HHSN27200008, unpublished data. Mice were given irradiated food and sterile water containing 50 μg/ml baytril (Bayer) ad libitum. 100 µl of purified mucoricin (0.1 mg/ml) was then injected into the tail vein on days 0, +2, and +4. The differences in survival between normal mice receiving vehicle (*i.e.* PBS) and those received toxin were compared by the Log Rank test. The primary efficacy endpoint was time to morbidity.

Mouse tissues were fixed in 10% ZnCl_2_ formalin solution prior to histopathological examination. The fixed organs were dehydrated in graded alcohol solutions, embedded in paraffin, and 5-μm sections were cut and stained with H&E^77^. Cumulative histopathological scores of hemorrhages, neutrophil infiltration (inflammation), and edema were used to determine the effects of toxin by observing 5 fields per slide. The observer was not told the origin of the samples.

### Mucoricin RNAi knockdown

RNAi knockdown of mucoricin was employed using our previously described RNAi method^20^. Briefly, a 330-bp mucoricin transcript was PCR amplified using 5’-AAATTTAAAA<u>GCATGC</u>ACACACAAAAGTATGAAGATTGCT-3’ and 5’-CTGCTTACCAT<u>GGCGCGCC</u>CAAATGGCACTAATTCCCAGC-3’ primers and cloned into the *Sph*I and *Asc*I sites of pRNAi-pdc^78^. The inverted repeat fragment was PCR amplified by 5’-TTAAGCGATC<u>GCTAGCAC</u>ACACAAAAGTATGAAGATTGCT-3’ and 5’-TTATTCTTATAGC<u>CCGCGG</u>CAAATGGCACTAATTCCCAGC-3’ at cloned downstream the intro fragment at the *Nhe*I and *Sac*II sites. The developed construct was transformed into *R. delemar* pyrF mutant (strain M16)^55^ using the biolistic delivery system (BioRad), and the homogenous transformants were selected on minimal medium lacking uracil^11^. The down regulation of mucoricin expression was confirmed by qRT-PCR using primers 5’-CTTGGATATCCGTGGAGGTGA-3’ and 5’-GGCAGCTTCTTCGACCATCT-3’ as described before^12^ and by confocal microscopy using immunostaining (see below)^11^.

### Secretion/shedding of mucoricin into the culture supernatant

Wild-type *R. delemar* spores (2 x 10^4^/100 μl/well), *R. delemar* transformed with the empty plasmid or those transformed with mucoricin RNAi were grown in 96-well plates for 24 h at 37℃ followed by additional 24 h of incubation in the presence or absence of 2-fold serially diluted amphotericin B (0.06-32 μg/ml). 100 μl of culture supernatant samples from each well were collected and stored at −20℃ until used for toxin detection by ELISA. To determine corresponding fungal growth, 100 μl/well XTT substrate (0.20 mg/ml activated with 6.25 μM menadione) was added to the remaining *R. delemar* culture plate.^79^ After a 2 h incubation at 37℃, absorbance at 450 nm was measured for metabolized XTT. Sandwich ELISAs were used to detect and quantify mucoricin in the cell-free supernatants. Briefly, 96-well plates were coated with 2 μg/ml mouse anti-*R. delemar* toxin monoclonal antibodies at 4℃ overnight. After washing the plate with 1X PBST (PBS+ 0.05% Tween-20) 5 times, diluted recombinant mucoricin or undiluted culture supernatant samples were added to the ELISA plate. Bound mucoricin was detected by the IgG anti-toxin antibodies (2 μg/ml), and subsequently by HRP-IgG anti-rabbit IgG (Jackson ImmunoResearch, product number 111-035-144) and a TMB substrate detection system (Invitrogen). A standard curve was generated using linear regression of OD_450_ of known recombinant mucoricin concentrations and the concentrations of toxin in the medium were extrapolated from the standard curve.

### Confocal microscopy

IgG anti-toxin was used to localize the toxin in the *Rhizopus* fungus^10^. Fungal spores (10^5^/ml) were pre-germinated in YPD media at 1, 4, or 12 h. Each fungal stage was fixed in 4% paraformaldehyde followed by permeabilization for 10 min in 0.1% Triton X-100. The permeabilized fungal growth stages were incubated with the IgG anti-toxin for 2 h at room temperature. The fungal stages were then washed 3 times with Tris-buffered saline (TBS, 0.01 M Tris HCl [pH 7.4], 0.15 M NaCl) containing 0.05% Tween 20 and counterstained with anti-rabbit IgG Alexa Fluor 488 (Life Technologies, Cat # A-11034). The stained fungi were imaged with Leica confocal microscope at excitation wavelength of 488 nm. The final confocal images were produced by combining optical sections taken through the z axis.

### *In vivo* virulence studies and immunohistochemistry

Male ICR mice (≥20 g) were rendered DKA with a single intraperitoneal injection of 210 mg/kg streptozotocin in 0.2 ml citrate buffer 10 days prior to fungal challenge. On days –2 and +3 relative to infection, mice were given a dose of cortisone acetate (250 mg/kg). Diabetic ketoacidotic (DKA) mice were given irradiated food and sterile water containing 50 μg/ml Baytril (Bayer) ad libitum. DKA mice were infected intratracheally with fungal spores with a target inoculum of 2.5 × 10^5^ spores of RNAi-empty plasmid (Control strain) or RNAi-mucoricin (targeting mucoricin gene expression) in 25 μl. To confirm the fidelity of the inoculum, three mice were sacrificed immediately after inoculation, their lungs were homogenized in PBS and quantitatively cultured on PDA plates containing 0.1% triton, and colonies were counted after a 24-hour incubation period at 37°C. Average inhaled inoculum for RNAi-empty plasmid and RNAi-mucoricin were 8.6 x 10^3^ and 3.3 x 10^3^ spores from two experiments, respectively. Primary endpoint was time to moribundity analyzed by Kaplan Meier plots. In another experiment, DKA mice were infected as above and then sacrificed on Day +4 relative to infection, when their lungs and brains (primary and secondary target organs) were collected and processed for determination of tissue fungal burden by qPCR^21^. The ability of the IgG anti-toxin to protect against *Rhizopus* infection was also evaluated in the DKA mouse model. Briefly, DKA mice were infected with *R. delemar* 99-880 as above (average inhaled inoculum of 5.6 x 10^3^ spores from two experiments) and 24 h later were injected intraperitoneally with either a 30 μg of IgG anti-toxin or normal rabbit IgG (R & D Systems, Cat # AB-105-C). The survival of mouse and tissue fungal burden of target organs collected on Day +4 post infection served as endpoints as above. Furthermore, histopathological examination was carried out on sections of the organs harvested on Day +4 post infection. These organs were fixed in 10% zinc formalin and processed as above for histological examination with H&E, PAS or Grocott staining.

Apoptotic cells in the lung were detected by immunohistochemistry using the ApopTag *in situ* apoptosis detection kit (EMD Millipore) following the manufacturer’s directions. Briefly, paraffin-embedded sections were rehydrated in Histo-Clear II (National Diagnostics) and alcohols followed by washing with phosphate-buffered saline (PBS). The sections were pre-treated with 20 μg/ml Proteinase K (Ambion) in PBS for 15 min at room temperature. Endogenous peroxidases were blocked by incubation of the slides for 15 min in 3% hydrogen peroxide. Sections were incubated with equilibration buffer (EMD Millipore) for 30 sec at RT, followed by terminal deoxynucleotidyl transferase (TdT; EMD Millipore) at 37°C for 1 h. Sections were further exposed to anti-Digoxignenin for 30 min at RT, and the positive reaction was visualized with DAB 3, 3-diaminobenzidine (DAB) substrate (Thermo Scientific). After counterstaining the specimens with 0.5% methyl green (Sigma), they were imaged by bright field microscopy. For quantification, apoptotic areas were quantified using PROGRES GRYPHAX^®^ software (Jenoptik).

### Immunofluorescence staining for mucoricin in human tissue samples

Paraffin-embedded human lung tissue from a patient diagnosed with disseminated mucormycosis^9^ or a patient with proven invasive pulmonary aspergillosis were cut into 5 µm sections that were then mounted onto glass slides. Organ sections on slides were deparaffinized and rehydrated with an ethanol gradient (100%-70%) followed by incubation of the slides in water and heat-induced antigen retrieval in sodium citrate buffer (10 mM, pH 6). Sections were blocked with 3% bovine serum albumin (BSA) in PBS (BSA-PBS), incubated for 1 h with 1:50 dilution of the IgG anti-mucoricin in PBS, washed twice in PBS, stained with 1:500 dilution of the appropriate goat anti-rabbit IgG Alexa Fluor^®^ 488 (Life Technologies, Cat #A-11034) in 1x PBS, followed by DNA staining with 1 µM TOPRO-3 iodide (642/661; Invitrogen) and staining of the fungal hyphae with 100 μg/ml Fluorescent Brightener 28 (Sigma-Aldrich, Cat #475300). After washing with 1x PBS, slides were mounted in Prolong Gold antifade media (Molecular Probes). Images were acquired using a laser-scanning spectral confocal microscope (TCS SP8; Leica), LCS Lite software (Leica), and a 40× Apochromat 1.25 NA oil objective using identical gain settings. A low fluorescence immersion oil (11513859; Leica) was used, and imaging was performed at room temperature. Serial confocal sections at 0.5 μm steps within a z-stack spanning a total thickness of 10 to 12 μm of tissue, and 3D images were generated using the LCS Lite software. Corresponding tissue sections from the same area were also stained with hematoxylin and eosin.

### Statistical analysis

The data was collected and graphed and statistically analyzed using Microsoft Office 360 and Graph Pad 8.0 for Windows or Mac (GraphPad Software, La Jolla, CA, USA). Cell damage and gene expression were analyzed using one-way analysis of variance (ANOVA) using Dunnett’s Multiple Comparison Test. The non-parametric log-rank test was used to determine differences in mouse survival times. Differences in tissue fungal burdens were compared by the non-parametric Wilcoxon rank sum test for multiple comparisons. *P* < 0.05 was considered as significant. All *in vitro* experiments were performed at least in triplicate and replicated at least once.

### Study approval

All procedures involving mice were approved by the IACUC of The Lundquist Institute for Biomedical Innovations at Harbor-UCLA Medical Center, according to the NIH guidelines for animal housing and care. Human endothelial cell collection was approved by the IRB of The Lundquist Institute for Biomedical Innovations at Harbor-UCLA Medical Center. Because umbilical cords are collected without donor identifiers, the IRB considers them medical waste not subject to informed consent. The purification and testing of ricin were approved by the IRB at UT Southwestern and carried out under BSL3 guidelines. Approval for the collection of tissue samples from the patients with mucormycosis and invasive pulmonary aspergillosis was obtained and the Ethics Committee of the University Hospital of Heraklion, Crete, Greece (5159/2014). Τhe patients provided written informed consent in accordance with the Declaration of Helsinki.

### Data availability

Source data are provided with this paper.

## Extended Data Figures Legends

**Extended Data Figure 1.**
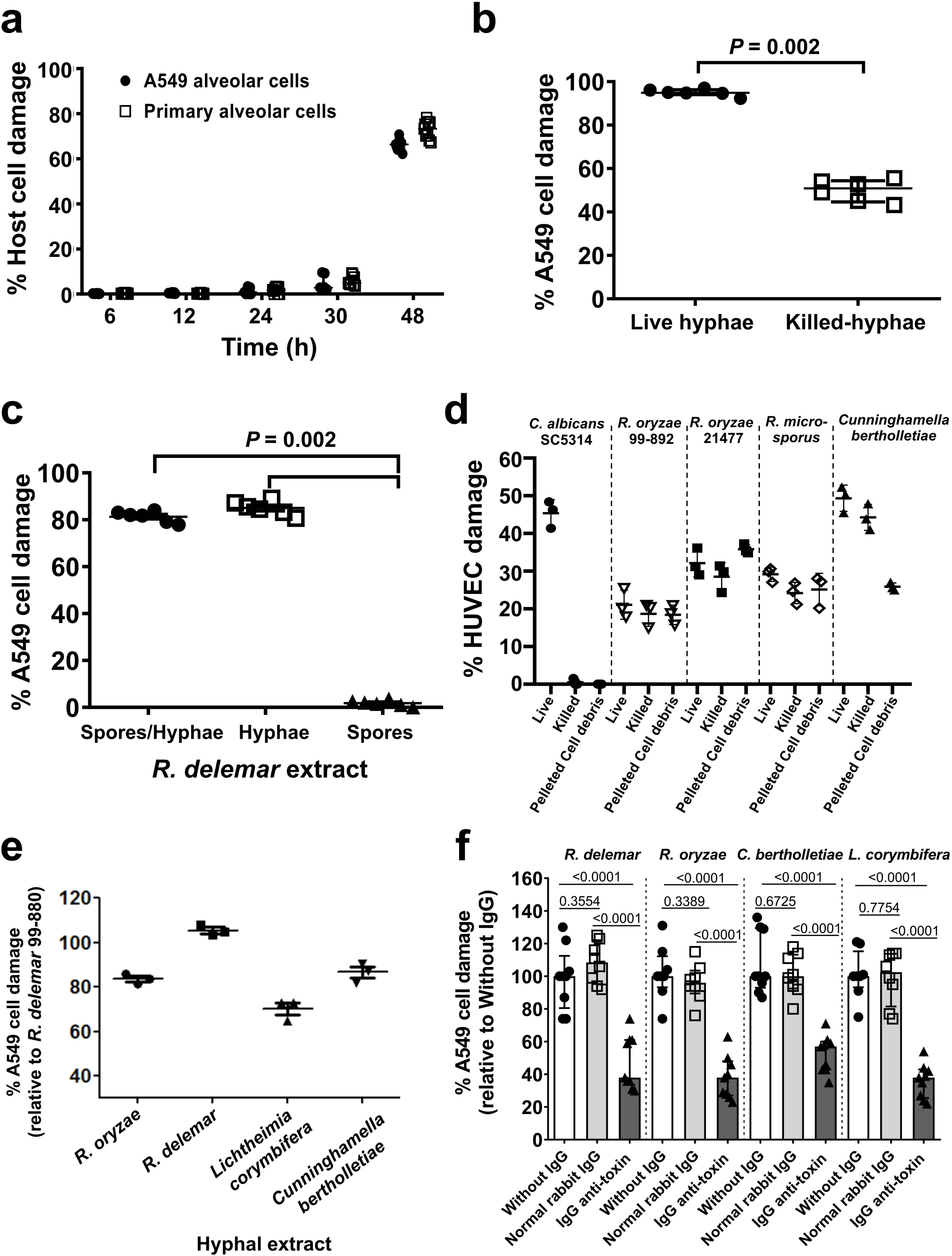
A heat stable and hyphae-associated Mucorales extract damages mammalian host cells *in vitro*. **(a)** *R. delemar* caused time dependent alveolar epithelial cell damage (n=9 wells/time point, pooled from three independent experiments). Data are median ± interquartile range. **(b)** Heat-killed *R. delemar* hyphae showed ∼50% damage to mammalian cells compared to ∼100% damage caused by living hyphae (n=6 wells/group, pooled from three independent experiments). Data are median ± interquartile range. Statistical analysis was performed by using Mann-Whitney non-parametric (two-tailed) test comparing live vs killed hyphae. **(c)** Extracts from comparable wet weight of *R. delemar* hyphae/spores, or hyphae, but not spores, damaged alveolar epithelial cells (n=6 wells/group, pooled from three independent experiments). Data are median ± interquartile range. Statistical analysis was performed by using Mann-Whitney non-parametric (two-tailed) test comparing spores vs spore/hyphae or hyphae. **(d)** Disrupted pellet from Mucorales germlings containing the cell-associated fraction was compared to live or heat-killed cells in causing injury to HUVECs (n= 3 wells/group, pooled from three independent experiments). Data are median ± interquartile range. **(e)** Fungal hyphae from representative clinical Mucorales isolates ground in liquid nitrogen and extracted with mammalian cell culture caused significant A549 alveolar epithelial cell damage (n= 3 wells/Mucorales, pooled from three independent experiments). Data are median ± interquartile range. **(f)** IgG anti-*R. delemar* toxin but not normal rabbit IgG (50 μg/ml) blocked host cell damage caused by heat-killed hyphae from different Mucorales (n=8 or 9 replicates/treatment/Mucorales, pooled from three independent experiments). Data presented as median ± interquartile range. Statistical analysis was performed by Mann-Whitney non-parametric (two-tailed) test comparing IgG anti-toxin *vs.* without IgG or normal rabbit IgG.

**Extended Data Figure 2.**
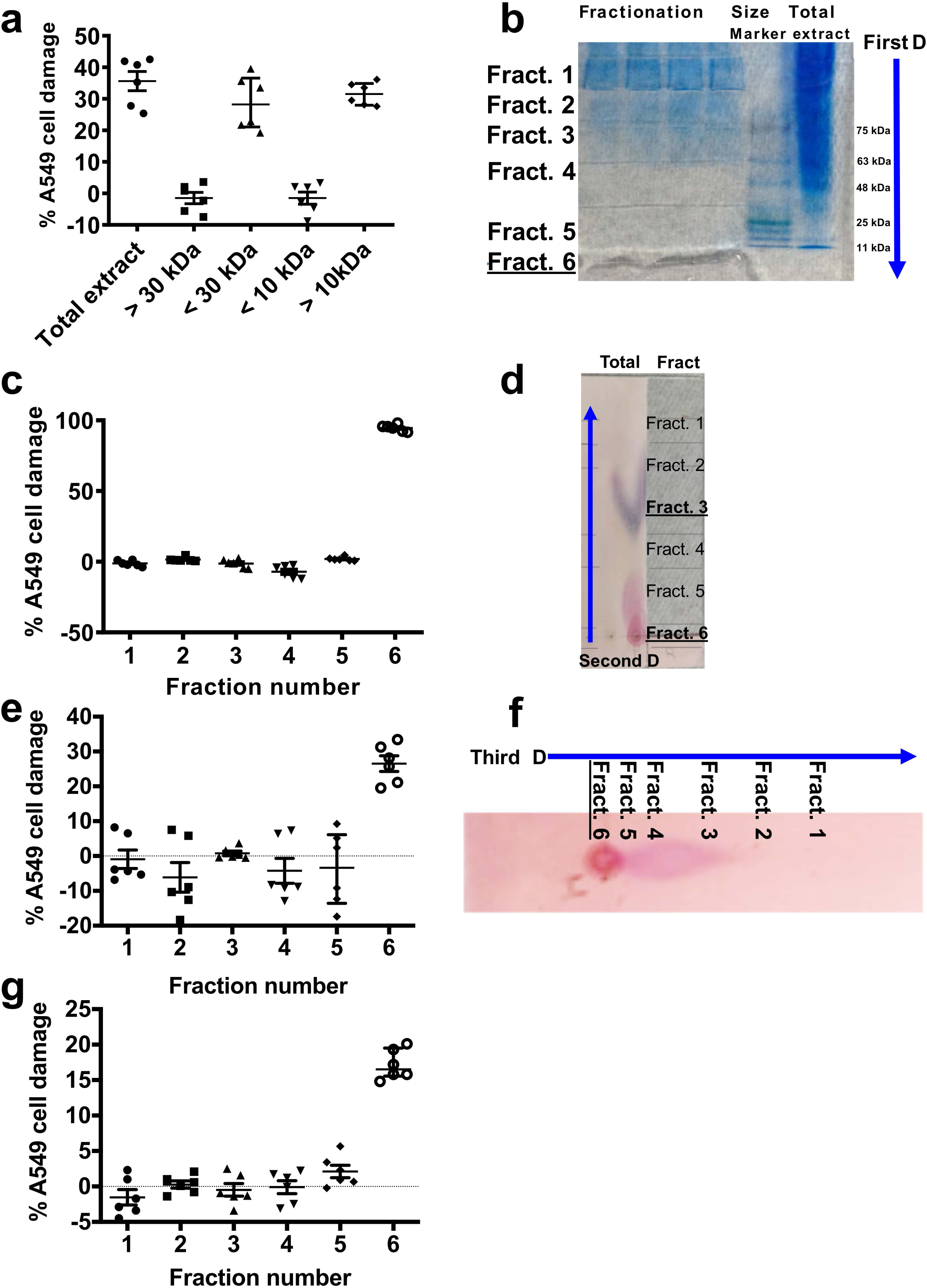
Fractionation and purification of *R. delemar* toxin. **(a)** Size exclusion of hyphae extracts indicating a 10-30 kDa fraction causing A549 cell damage (n=6 wells/fraction, pooled from three independent experiments). Data are median ± interquartile range. **(b)** Native polyacrylamide fractionation of hyphae extract and **(c)** its corresponding A549 cell damage, showing fraction # 6 causing injury. (n=6 wells/fraction, pooled from three independent experiments). Data are median ± interquartile range. **(d)** Cellulose plate separation of fraction # 6 purified from the polyacrylamide gel and **(e)** its corresponding A 549 cell damage, showing a high polar fraction #6 causing injury. Data are n=6 wells/fraction, and pooled from three independent experiments. Data are median ± interquartile range. **(f)** Third dimension fractionation of the previous fraction # 6 on cellulose plates and **(g)** its corresponding A549 cell injury (n=6 wells/fraction, pooled from three independent experiments). Data are median ± interquartile range.

**Extended Data Figure 3.**
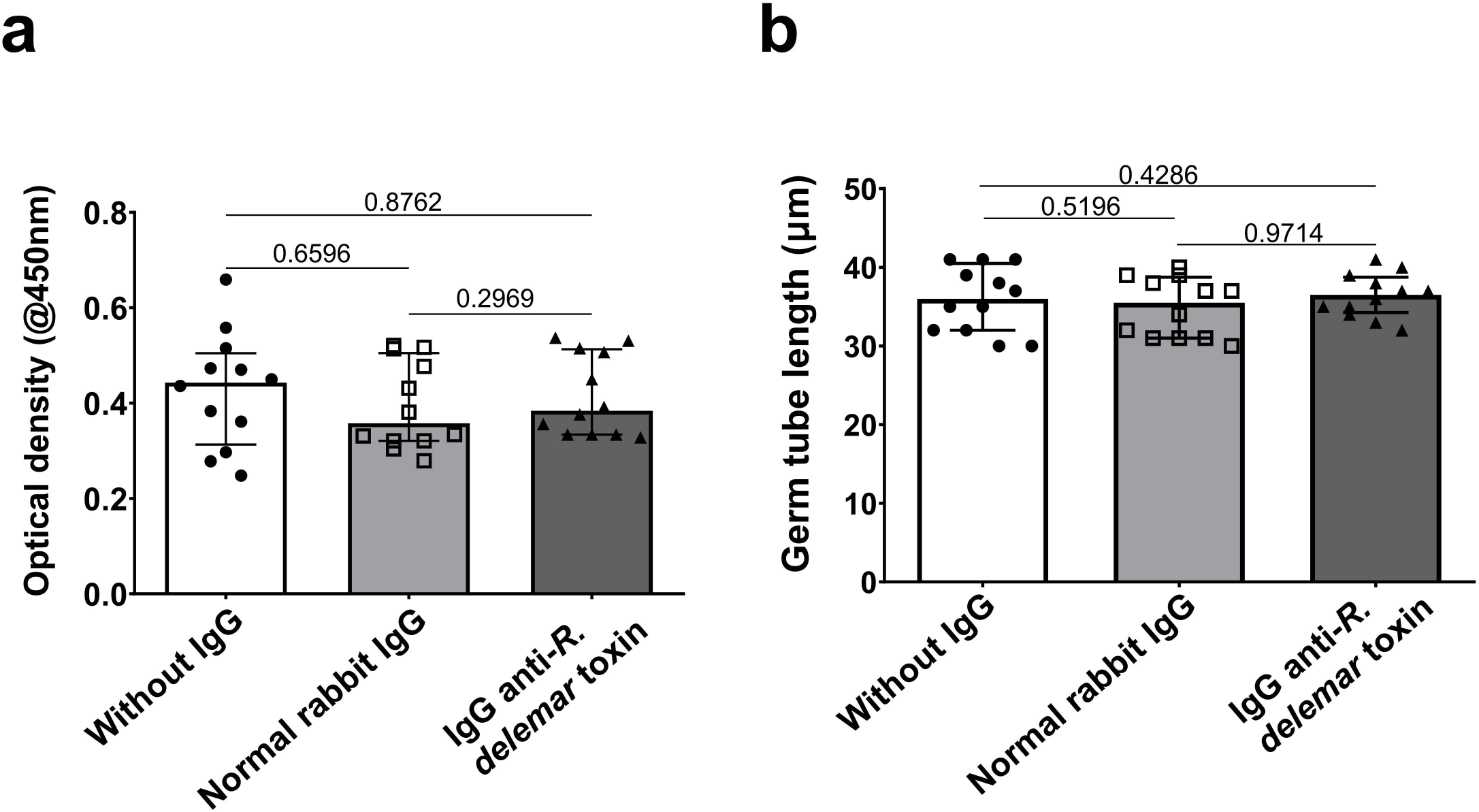
IgG anti-toxin had no effect on growth or germination of *R. delemar.* **(a)** Fungal spores (10^4^/ml) were inoculated in 96-well plates with or without 50 μg/ml IgG anti-toxin or normal rabbit IgG for 6 h prior to measuring absorbance at 450 nm. (n=12 wells, data pooled from three independent experiments) Data presented as median + interquartile range. Statistical analysis was performed by Mann-Whitney non-parametric (two-tailed). (**b**) *R. delemar* spores (10^4^/ml) were germinated at 37°C for 6 h prior to measuring the germ tube length using light microscopy equipped with a micometer lens. Each data point represents 20-50 germ tubes/HPF. (n=12 wells, data pooled from three independent experiments) Data presented as median + interquartile range from three experiments. Statistical analysis was performed by Mann-Whitney non-parametric (two-tailed).

**Extended Data Figure 4.**
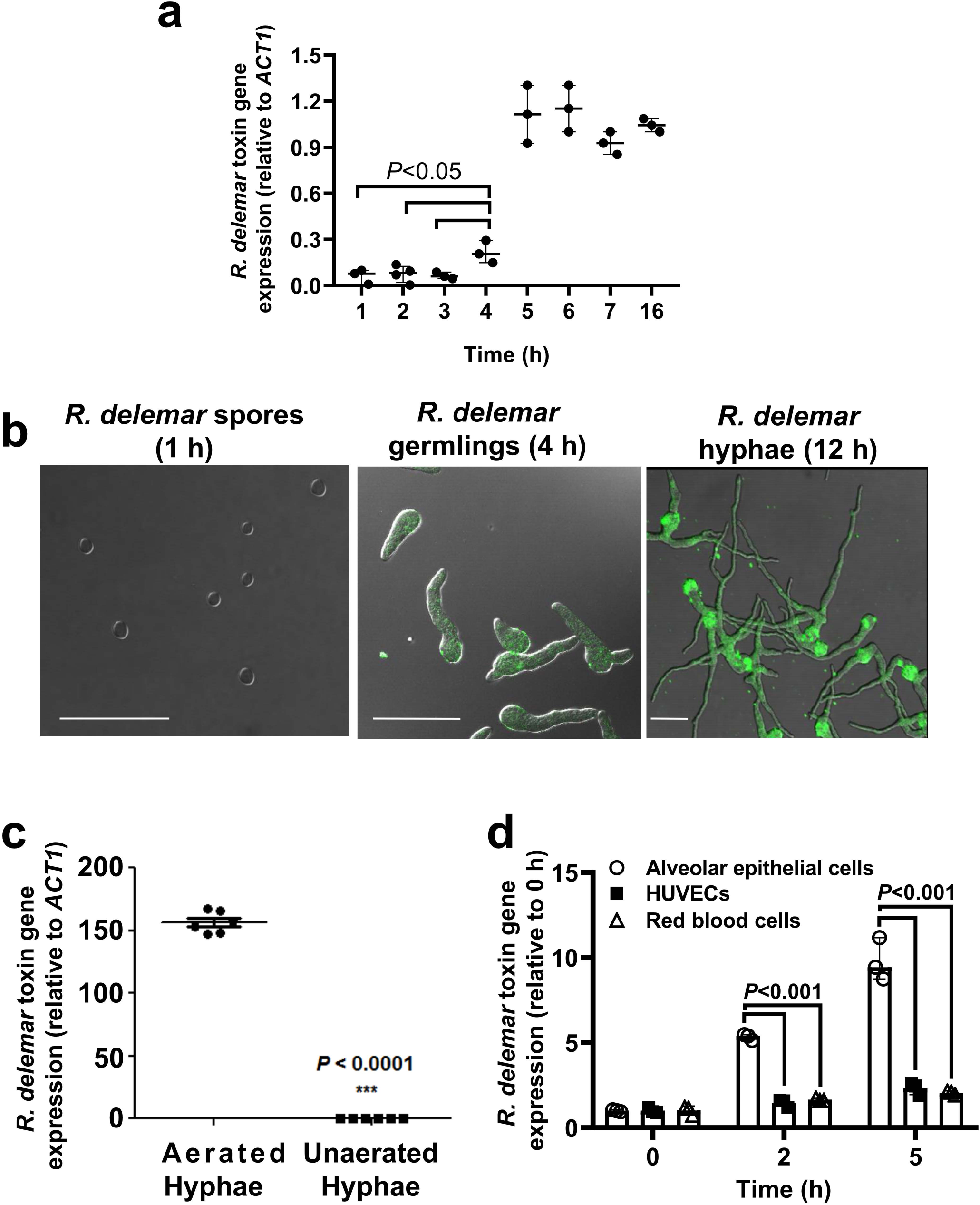
Putative toxin gene expression is cell-, time- and oxygen-dependent. **(a)** Toxin gene expression in *R. delemar* germinating cells in YPD medium. Data (n=3 wells/timepoint, pooled from three independent experiments) are presented as median ± interquartile range. Statistical analysis was performed by using unpaired t-test (two-tailed). **(b)** Confocal imaging of Alexa Flour 488-labelled IgG anti-toxin (green) during the growth of *R. delemar* from spores to hyphae. Scale bar is 50 µm. **(c)** Toxin gene expression from *R. delemar* hyphae grown in YPD culture in sufficient versus limited oxygen (n=6 wells, data pooled from three independent experiments). Data presented as median ± interquartile range. Statistical analysis was performed by using unpaired t-test (two-tailed). **(d)** Toxin gene expression analysis of fungal germlings on different cell types showed a time dependent expression on alveolar epithelial cells compared to HUVECs and erythrocytes (n=3 wells/group, pooled from three independent experiments). Data presented as median ± interquartile range. Statistical analysis was performed by using unpaired t-test (two-tailed).

**Extended Data Figure 5.**
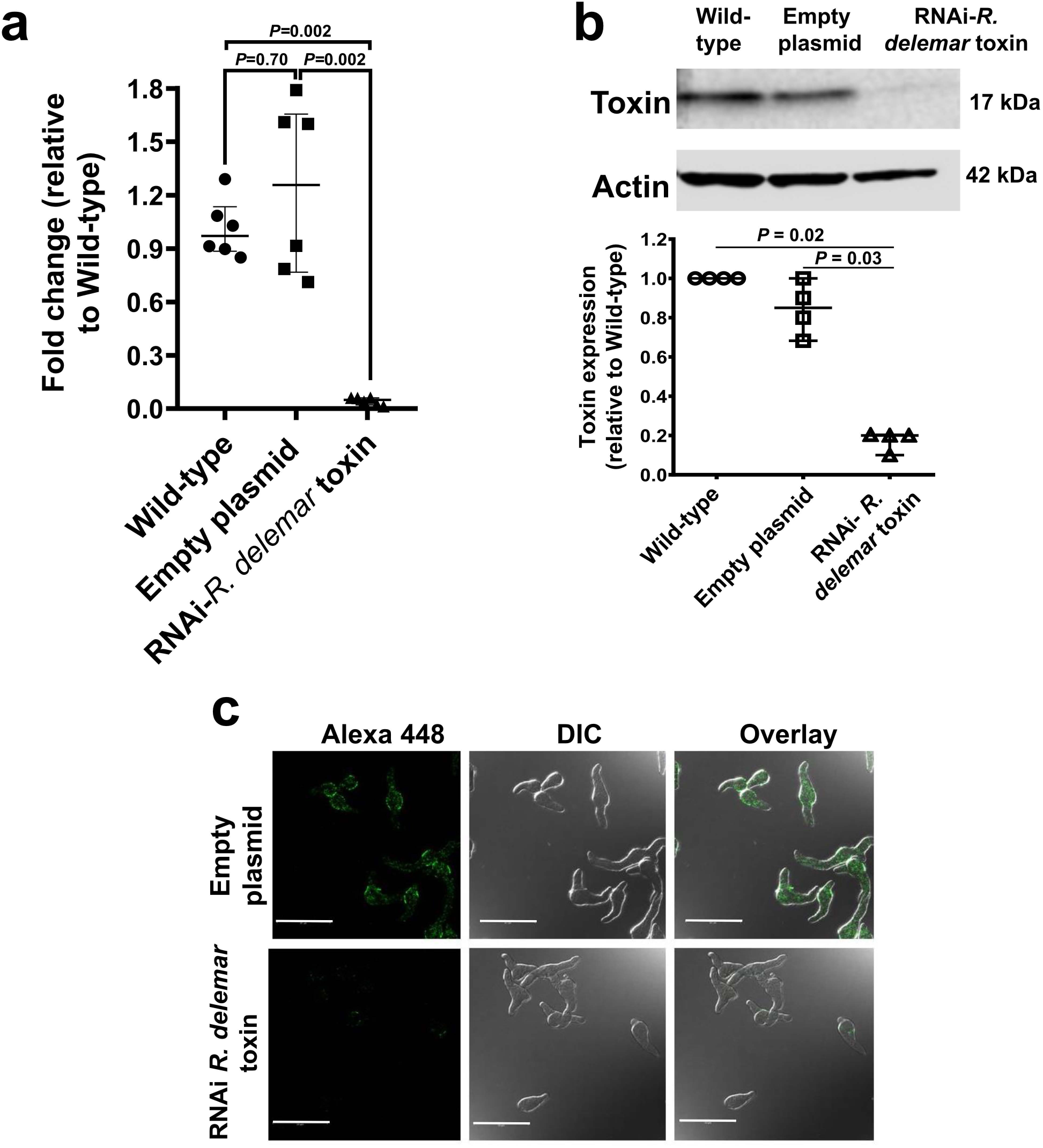
RNAi targeting the putative *R. delemar* toxin inhibits its expression. **(a)** *R. delemar* spores were transformed with RNAi plasmids targeting the putative toxin (RNAi-toxin) or empty plasmid (Empty-plasmid) using biolistic delivery system. Cells were grown in minimal medium without uracil for 24 h prior to extracting RNA (n=6/group, pooled from three independent experiments). Data presented as median ± interquartile range. Statistical analysis was performed by using Mann-Whitney non-parametric (two-tailed) test comparing RNAi- *R. delemar* toxin vs wild-type or empty plasmid **(b)** Representative Western blot and densitometry analyses of the wild-type, empty plasmid, or RNAi toxin strains (n=4 pictures data pooled from four independent experiments) Data presented as median ± interquartile range. Statistical analysis was performed by using Mann-Whitney non-parametric (two-tailed) test comparing RNAi- *R. delemar* toxin *vs.* wild-type or empty plasmid. **(c)** confocal images showing reduced expression of the toxin in the RNAi toxin mutant. Scale bar is 50 µm.

**Extended Data Figure 6.**
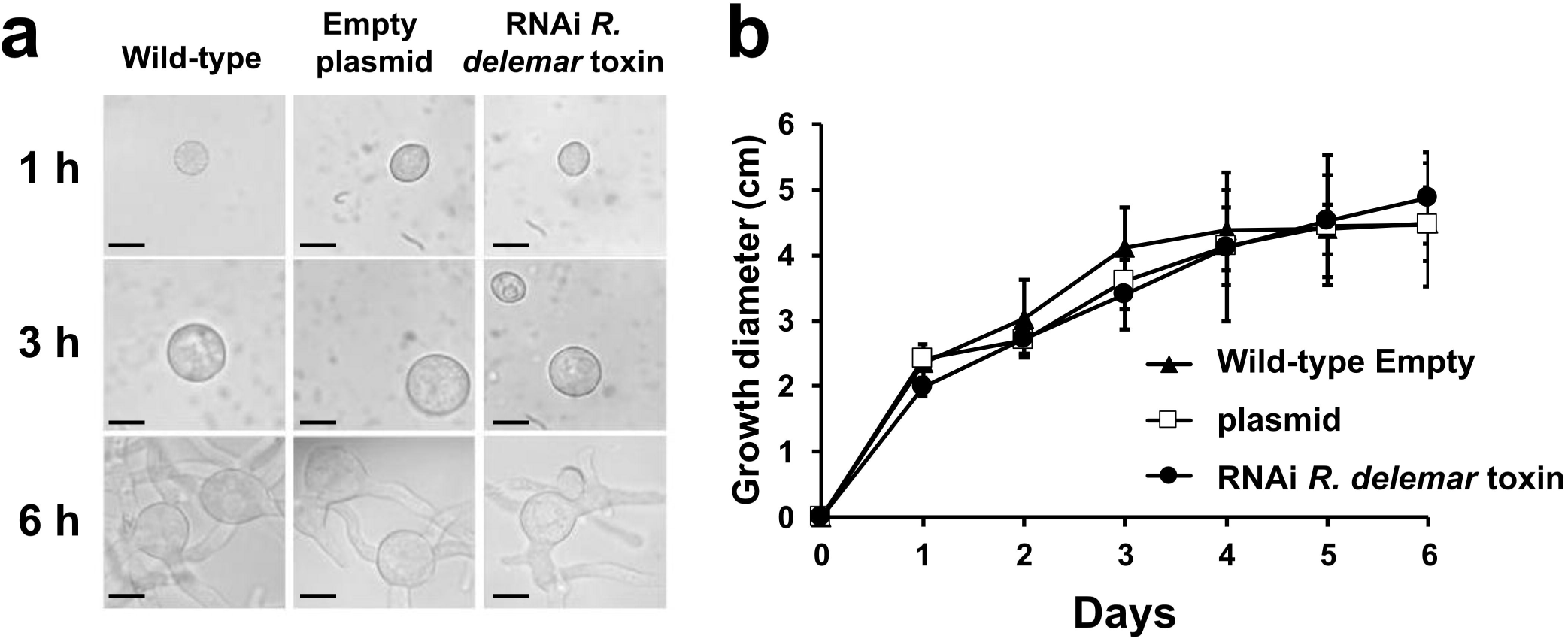
Down regulation of *R. delemar* toxin by RNAi did not affect germination or the growth of the fungus. **(a)** Wild-type *R. delemar,* RNAi empty plasmid, or RNAi toxin strains were germinated in minimal medium without uracil at 37°C with shaking. At times, samples were taken from the medium and examined by light microscopy. Scale bar is 5 µm. (**b**) 10^5^ spores of wild-type *R. delemar,* RNAi empty plasmid, or RNAi toxin strains were plated in the middle of the minimal medium without uracil agar plates for several days at 37°C and the colony diameter measured (n=6 plates/group, pooled from three independent experiments). Data are presented as median ± interquartile range.

**Extended Data Figure 7.**
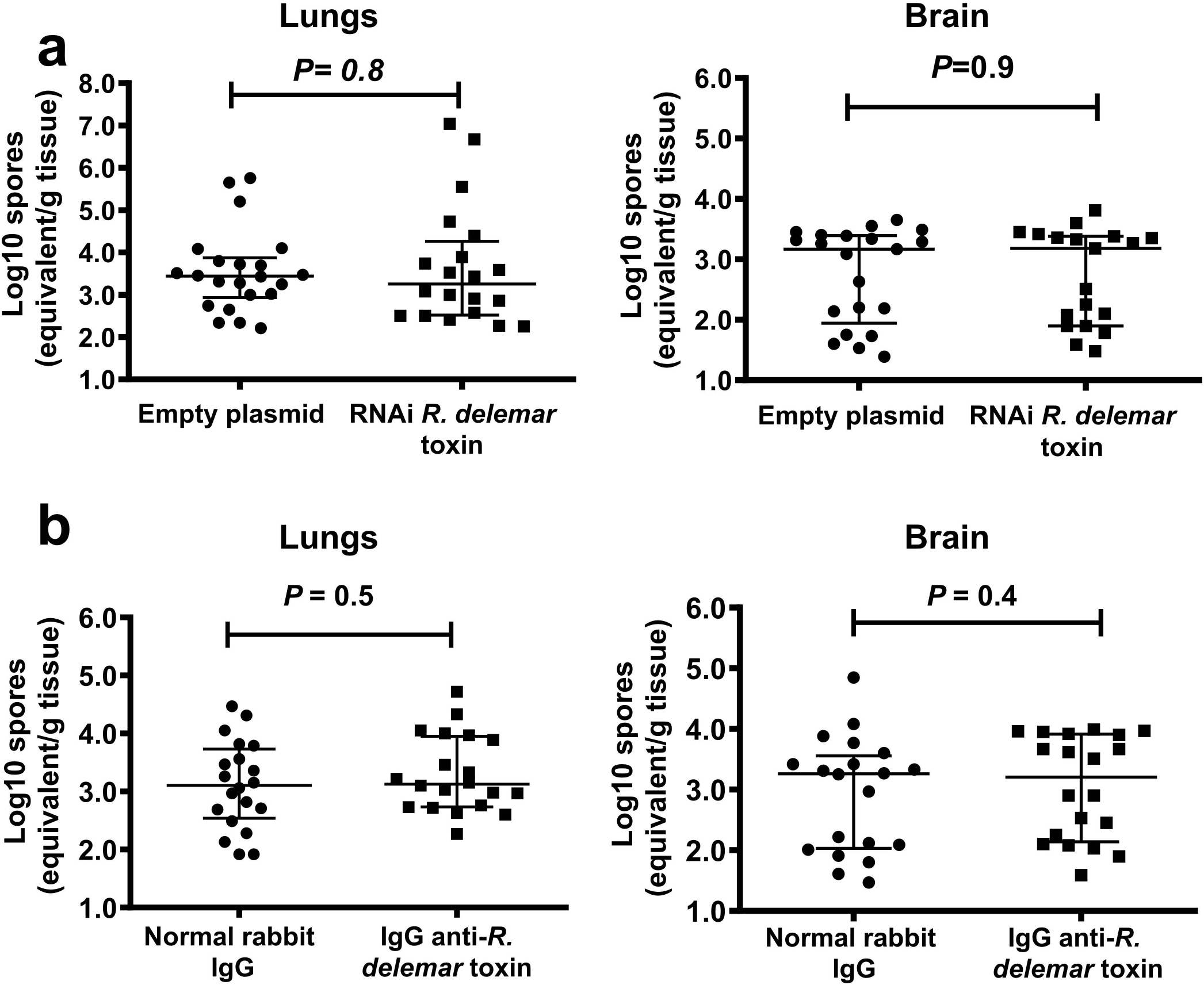
Effect of blocking the expression or the function of *R. delemar* toxin on fungal burdens in mice. **(a)** Inhibition of the toxin by RNAi did not affect the fungal burden in the lungs or brain of mice harvested on Day +4 post infection (average inoculum from two experiments of 1.4 x 10^4^ for empty plasmid [n=22 mice] *vs.* 1.3 x 10^4^ for RNAi toxin mutants [n=20 mice]). Data are pooled from two independent experiments and presented as median ± interquartile range. Statistical analysis was performed by using Mann-Whitney non-parametric (two-tailed) test comparing RNAi-*R.delemar* toxin *vs.* Empty plasmid. **(b)** The IgG anti-*R. delemar* toxin had no effect on the fungal burden of lungs or brains of DKA mice harvested on Day +4 post intratracheal infection with wild-type *R. delemar* (average inhaled inoculum of 5.6 x 10^3^ spores from two experiments [n=20 mice]). Data are pooled from two independent experiments and presented as median ± interquartile range). Statistical analysis was performed by using Mann-Whitney non-parametric (two-tailed) test comparing IgG anti-*R.delemar* toxin *vs.* normal rabbit IgG.

**Extended Data Figure 8.**
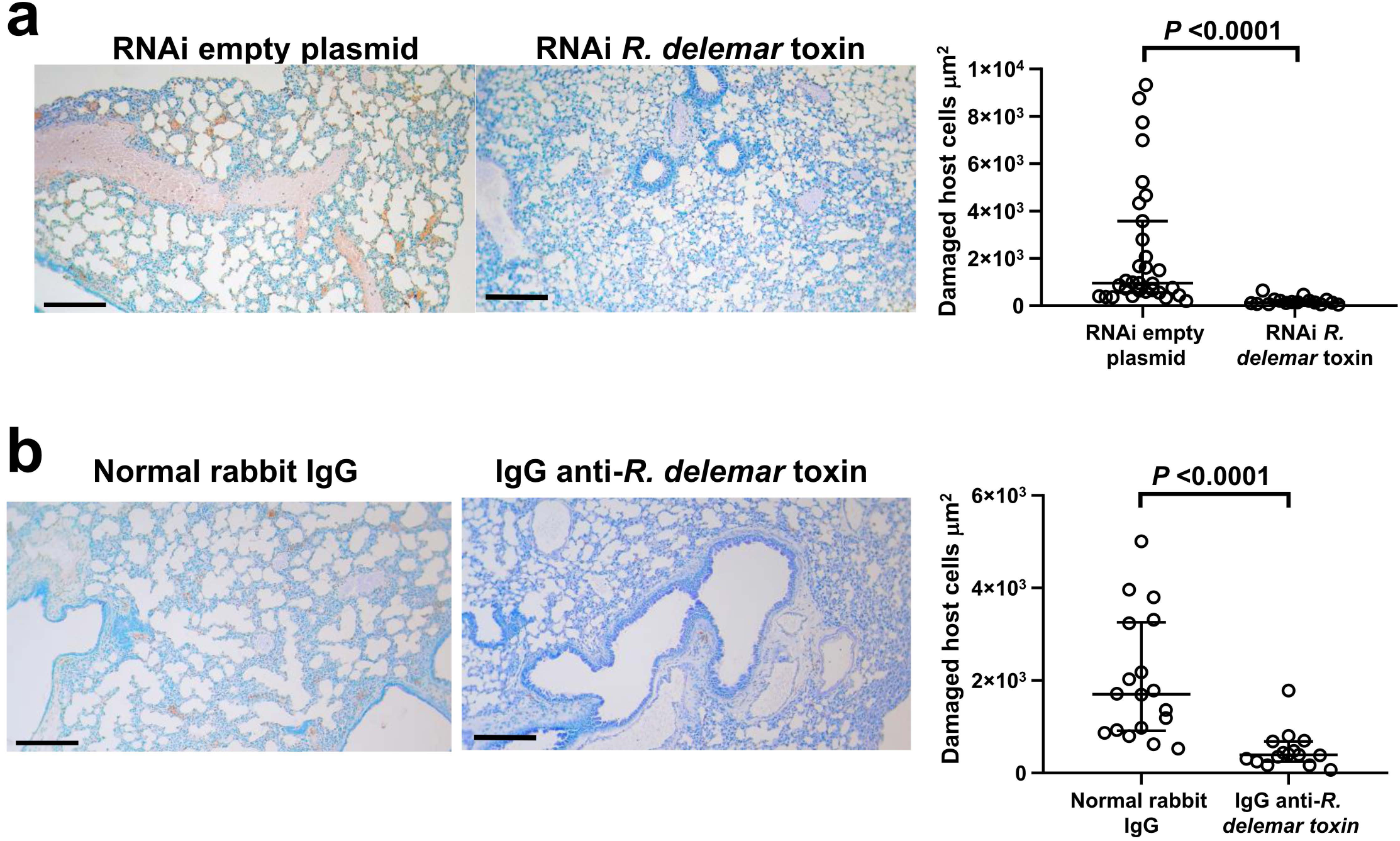
Histology of organs showing involvement of the toxin in tissue damage. **(a)** Damaged lung tissues (brown color) of mice infected with *R. delemar* transformed with RNAi empty plasmid (n=31 field counts) or RNAi toxin. Statistical analysis was performed by using Mann-Whitney non-parametric (two-tailed) test. Scale bar is 200 µm. (**b**) Damaged lung tissues from mice infected with wild-type *R. delemar* and treated with either normal rabbit IgG (n= 18 field counts) or IgG anti-toxin (n= 18 field counts) were quantified by ApopTag kit. Data were pooled from two independent experiments, are presented as median + interquartile range. Statistical analysis was performed by using Mann-Whitney non-parametric (two-tailed) test. Scale bar is 200 µm.

**Extended Data Figure 9.**
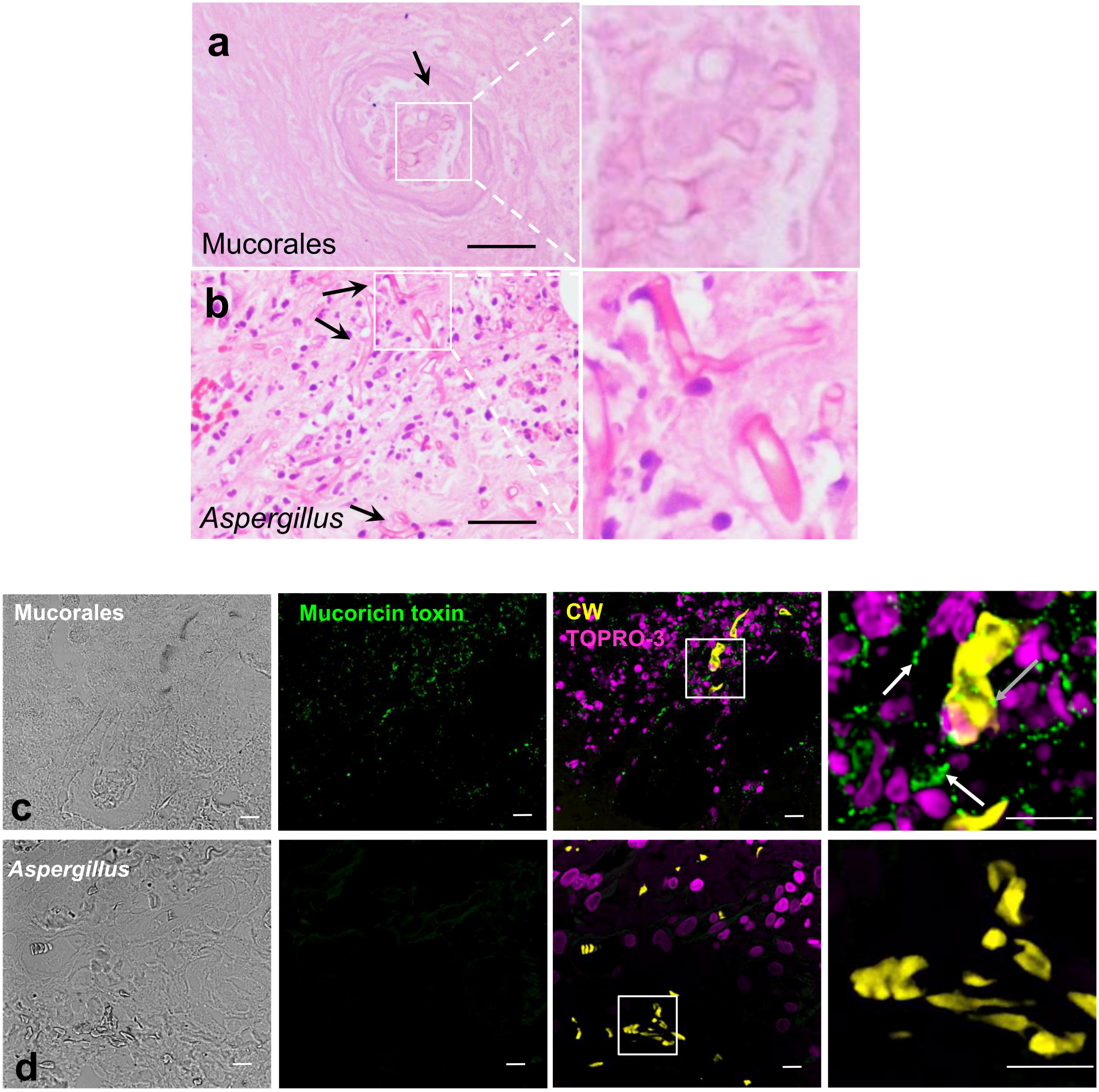
*R. delemar* toxin is expressed in lung tissue collected from a mucormycosis patient but not in lung samples from an aspergillosis patient. H&E staining of lung tissues from mucormycosis **(a)** or aspergillosis **(b)** patients showing broad aseptate hyphae with angioinvasion (Mucorales) and thinner septated hyphae of *Aspergillus.* Scale bar is 10 μm. Box magnification 1400 X. Staining of a mucormycosis **(c)** or aspergillosis **(d)** patient lungs using IgG anti-toxin (green color). Mucorales or *Aspergillus* hyphae are shown in yellow (stained with calcofluor white) and nuclei are shown in magenta. *R. delemar* toxin staining is shown in association with hyphae (grey arrow) and released in the tissue (white arrow). Scale bar is 10 μm in all micrographs.

**Extended Data Figure 10.**
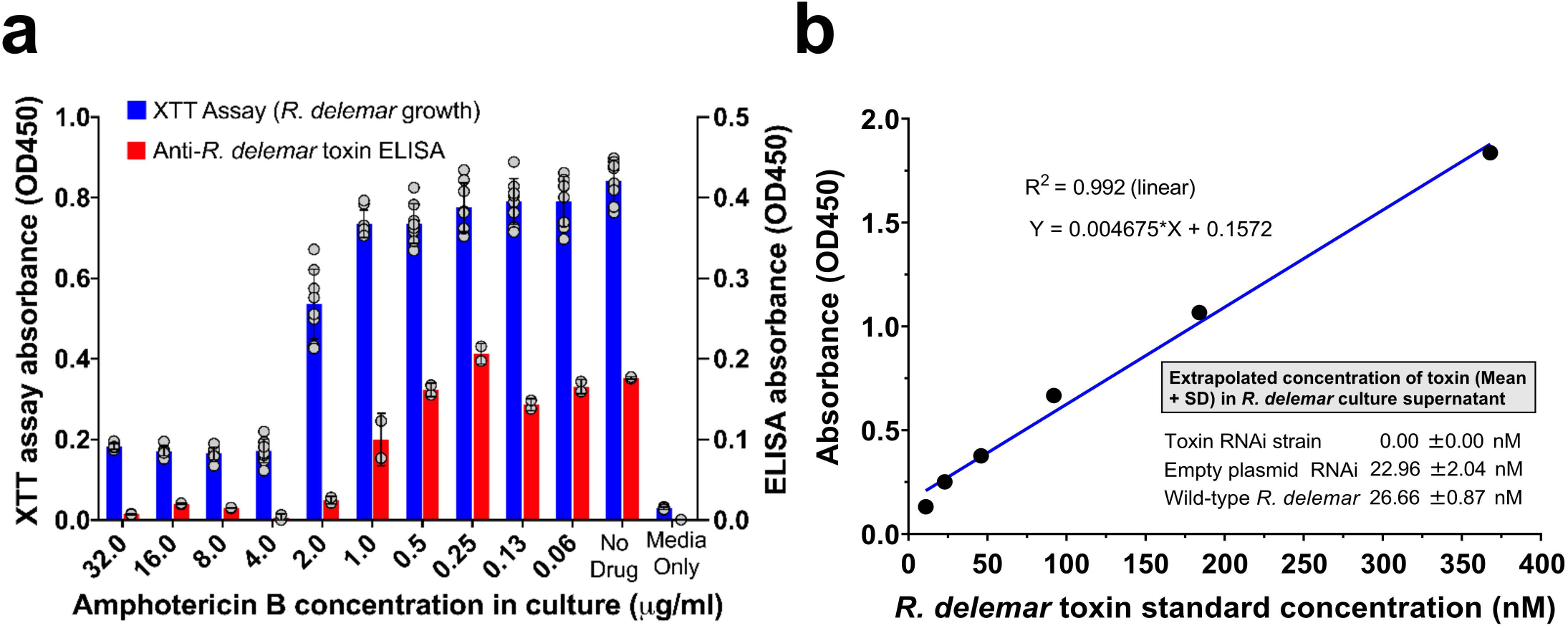
Secretion/shedding of *R. delemar* toxin in culture supernatant of growth media. **(a)** Cell-free culture supernatants were collected from *R. delemar* hyphae grown in the presence or absence of 2-fold dilutions of amphotericin B. The XTT assay was used to determine growth of *R. delemar* (left axis, blue bar, n=8 wells/amphotericin B concentration), while toxin release assayed by sandwich ELISA using anti-*R. delemar* mouse monoclonal IgG1 as the capture antibody and rabbit anti-*R. delemar* toxin IgG as the detector antibody (right axis, red bar, n=2 wells/amphotericin B concentration). Data in are representative of three independent experiments and presented as mean ± SD. **(b)** The released toxin concentration from *R. delemar* wild-type, *R. delemar* transformed with empty plasmid RNAi or *R. delemar* with RNAi-toxin was extrapolated from a standard curve using recombinant toxin in the same ELISA assay. Toxin concentrations (n= 3 samples from three independent experiments tested in duplicate in ELISA for each strain) are presented as mean ± SD.

## Supplementary Figure Legends

**Supplementary Figure 1. Incubation of lower inoculum of *R. delemar* with HUVECs induces minimal to no host cell injury.** Data are presented as % ^51^Cr-released from HUVECs challenged with 1 x 10^5^ spores of *R. delemar* for 5 hours after subtracting the amount of ^51^Cr-released from HUVECs without *R. delemar* challenge. (n= 10 data pooled from three independent experiments). Data presented as median + interquartile range.

**Supplementary Figure 2. CLUSTAL multiple sequence alignment by MUSCLE (3.8) between mucoricin and saporin from *Saponaria officinalis.*** The predicted Type 1 RIP domain in saporin (shown in yellow) aligned with sequence from mucoricin with 10 out of 17 amino acid residues conserved.

**Supplementary Table 1:** Results of BLAST search of a ricin-like toxin gene from *R. delemar* 99-880.

**Supplementary Table 2.** Ten proteins that are structurally similar to mucoricin. The 3-D model of mucoricin was used to identify structurally similar proteins in the protein data bank (PDB) by Tm align.

**Supplementary Table 3:** Ricin orthologues in different Mucorales and the presence of vascular leak and RIP motifs.

