## Supplementary Fig 1-2 and Table 2-3 for "Mucoricin is a Ricin-Like Toxin that is Critical for the Pathogenesis of Mucormycosis"

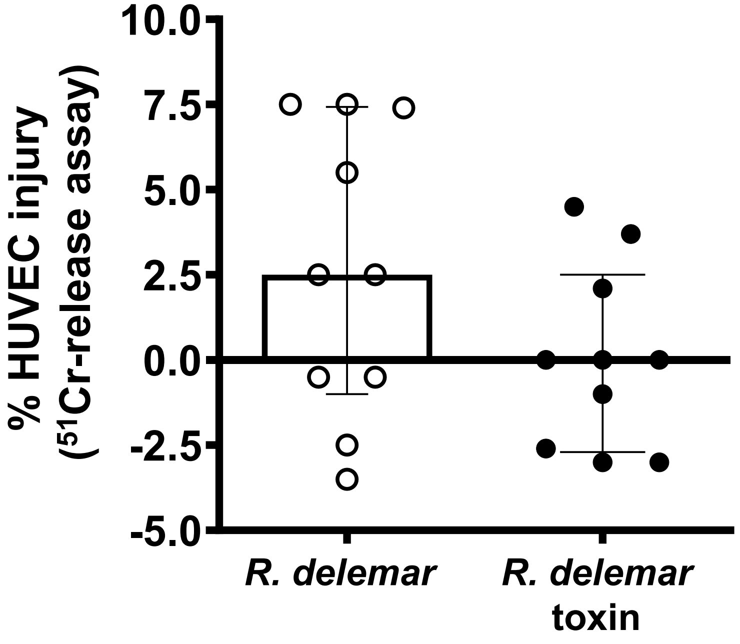


**Supplementary Fig. 1. Incubation of lower inoculum of *R. delemar* or its toxin with HUVECs induces minimal to no host cell injury.** Data are presented as % $^{51}$Cr-released from HUVECs challenged with 1 x $10^5$ spores of *R. delemar* or 50 µg/ml of *R. delemar* toxin for 5 hours. Data (n=10/group) are as median ± interquartile range from three experiments.

### Supplementary Fig. 2

```
saporin    VTSITLDLVNPTAGQYSSFVDKIRNNVKDPNLKYGGTDIAVIGPPSKEKFLRINFQSSRG
Mucoricin  -----------------MYFEE-----------------------GRLFFIKSQFNGRVL
                             :.::                       ..  *:. :*:.

saporin    TVSLGLKRDNLYVVAYLAMDNTNVNRAYYFRSEITSAESTALFPEATTANQKALEYTEDY
Mucoricin  DVEDGSTEDDANIIVYTQKYEDCLNQLW---------------------------RYENGY
            *. * . *:  ::.*    :  :*. :                           * :.*

saporin    QSIEKNAQITQGDQSRKELGLGIDLLSTSMEAVNKKARVVKDEARFLLIAIQMTAEAARF
Mucoricin  FINAKSAKV-----------------------LDIRGGEMQPESQIIQYAQKMVEEAANQ
               *.*::                       :: ..  :: *:.::  * :*. ***.

saporin    RYIQNLVIKNFPNKFNSENKVIQFEVNWKKISTAIYGDAKNGVFNKDYDFGFGKVRQVKD
Mucoricin  RWAID------------EDGYIFCEARPDLVLDIQGAEDEDCVPVILYERREGEVSA--N
           *:  :            *:  *  *.. . :     .: :: *    *:   *:*    :

saporin    LQMGLLMYLGKPK
Mucoricin  QRWELVPFEG---
```

**Supplementary Fig. 2. CLUSTAL multiple sequence alignment by MUSCLE (3.8) between mucoricin and saporin from *Saponaria officinalis.*** The predicted Type 1 RIP domain in saporin (shown in yellow) aligned with sequence from mucoricin with 10 out of 17 amino acid residues conserved.

**Supplementary Table 2. Ten proteins that are structurally similar to mucoricin.** The 3-D model of mucoricin was used to identify structurally similar proteins in the protein data bank (PDB) by Tm align.

| Rank | PDB Hit | Tm score | RMSD | Identity | Coverage | Protein Description | Classification | Link |
|---|---|---|---|---|---|---|---|---|
| 1 | 3ef2D | 0.947 | 1.05 | 0.166 | 0.986 | *Marasmius oreades* mushroom lectin (MOA) in complex with Gal-alpha(1,3)[Fuc-alpha(1,2)]Gal and Calcium | Sugar binding protein | https://www.rcsb.org/structure/3ef2 |
| 2 | 3vsfA | 0.871 | 1.28 | 0.248 | 0.912 | 1,3 Gal 43A, an exo-beta-1,3-Galactanase from *Clostridium thermocellum* | Sugar binding protein | https://www.rcsb.org/structure/3vsf |
| 3 | 3pg0A | 0.862 | 1.13 | 0.295 | 0.898 | 3-fold symmetric protein, Three Foil | De novo protein | https://www.rcsb.org/structure/3pg0 |
| 4 | 2x2tA | 0.838 | 1.55 | 0.157 | 0.912 | *Sclerotinia sclerotiorum* Agglutinin (SSA) in complex with Gal-beta1,3-Galnac | Cell adhesion | https://www.rcsb.org/structure/2x2t |
| 5 | 3nbeA | 0.835 | 1.6 | 0.164 | 0.912 | *Clitocybe nebularis* ricin B-like lectin (CNL) in complex with N,N'-diacetyllactosediamine | Sugar binding protein | https://www.rcsb.org/structure/3nbe |
| 6 | 4g9mA | 0.831 | 1.35 | 0.223 | 0.884 | Crystal structure of the *Rhizoctonia solani* agglutinin | Sugar binding protein | https://www.rcsb.org/structure/4g9m |
| 7 | 3phzA | 0.827 | 2.03 | 0.179 | 0.912 | *Polyporus squamosus* lectin bound to human-type influenza-binding epitope Neu5Aca2-6Galb1-4GlcNAc | Sugar binding protein | https://www.rcsb.org/structure/3phz |
| 8 | 5xg5A | 0.825 | 1.43 | 0.099 | 0.891 | Crystal structure of Mitsuba-1 with bound NAcGal | De novo protein | https://www.rcsb.org/structure/5xg5 |
| 9 | 2vseA3 | 0.825 | 1.95 | 0.187 | 0.905 | Structure and mode of action of a mosquitocidal holotoxin (*Lysinibacillus sphaericus*) | Toxin | https://www.rcsb.org/structure/2vse |
| 10 | 3a23B | 0.822 | 1.54 | 0.188 | 0.878 | Crystal Structure of beta-L-Arabinopyranosidase complexed with D-galactose | Hydrolase (glycosidase) | https://www.rcsb.org/structure/3a23 |

**Supplementary Table 3: Ricin orthologs in different Mucorales and the presence of vascular leak and RIP motifs**

| S. No. | S. No. | | MUSCLE Sequence Alignment (Percent Identity Matrix) | | | | | | | | | | | | | | | | | | | | Vacular Leak Motifs | | | | | Ribosome Inactivation Motifs (Reported in Ricin) | | | | | | | |
|---|---|---|---|---|---|---|---|---|---|---|---|---|---|---|---|---|---|---|---|---|---|---|---|---|---|---|---|---|---|---|---|---|---|---|---|
| | Organisms | | 1 | 2 | 3 | 4 | 5 | 6 | 7 | 8 | 9 | 10 | 11 | 12 | 13 | 14 | 15 | 16 | 17 | 18 | 19 | 20 | (x)D(y)-Motif [x= L, I, G or V], [y= V, L or S] | | | | | EAARF Motif | | | | WGRLS Motif | | | |
| | | | Ricinus communis | Choanephora cucurbitarum | Rhizopus delemar RA 99-880 | Mucor circinelloides f. circinelloides 1006PhL | Mucor circinelloides f. lusitanicus | Mucor circinelloides f. lusitanicus CBS 277.49 | Rhizopus azygosporus | Rhizopus microsporus | Rhizopus stolonifer | Mucor ambiguus | Rhizopus microsporus ATCC 52813 | Rhizopus microsporus var. microsporus | Phycomyces blakesleeanus NRRL1555 (-) | Absidia glauca | Absidia repens | Syncephalastrum racemosum | Hesseltinella vesiculosa | Lichtheimia corymbifera JMRC:FSU:9682 | Lichtheimia ramosa | Parasitella parasitica | | | | | | | | | | | | | |
| | Organisms' Name | Accession ID | NP_001310630.1 | OBZ89643.1 | EIE81863.1 | EPB83885.1 | KAF1807713.1 | OAD01005.1 | RCH82711.1 | ORE22231.1 | RCH89556.1 | GAN06585.1 | XP_023462404.1 | ORE05386.1 | XP_018295414.1 | SAL96273.1 | ORZ12009.1 | ORY99019.1 | ORX62783.1 | CDH57241.1 | CDS03748.1 | CEP13132.1 | LDV | IDV | IDS | GDL | VDV | Score | p-value | Q-value | Motif in Target Protein | Score | p-value | Q-value | Motif in Target Protein |
| 1 | Ricinus communis | NP_001310630.1 | 100 | 30 | 30 | 29 | 27 | 27 | 27 | 27 | 27 | 27 | 26 | 26 | 26 | 24 | 23 | 21 | 21 | 20 | 17 | 16 | Yes | | | | Yes | 19.81 | <0.00001 | 0.003 | EAARF | 20.92 /4.28 | <0.00001 / 0.0009 | 0.001/1 | WGRLS /RGRLT |
| 2 | Choanephora cucurbitarum | OBZ89643.1 | 30 | 100 | 81 | 78 | 76 | 76 | 85 | 75 | 75 | 75 | 74 | 74 | 80 | 45 | 45 | 46 | 45 | 47 | 43 | 73 | Yes | | | | | 4.25 | 0.0019 | 0.59 | EAANQ | | | | |
| 3 | Rhizopus delemar RA 99-880 | EIE81863.1 | 30 | 81 | 100 | 78 | 72 | 72 | 87 | 77 | 77 | 72 | 77 | 77 | 77 | 45 | 48 | 44 | 44 | 48 | 44 | 71 | Yes | | | | | 4.25 | 0.0019 | 0.59 | EAANQ | 4.28 | 0.0009 | 1 | EGRLF |
| 4 | Mucor circinelloides f. circinelloides 1006PhL | EPB83885.1 | 29 | 78 | 78 | 100 | 90 | 90 | 78 | 69 | 69 | 90 | 69 | 69 | 73 | 43 | 45 | 44 | 41 | 45 | 42 | 82 | Yes | | | | | 4.25 | 0.0019 | 0.59 | EAANQ | | | | |
| 5 | Mucor circinelloides f. lusitanicus | KAF1807713.1 | 27 | 76 | 72 | 90 | 100 | 100 | 73 | 66 | 66 | 97 | 68 | 68 | 73 | 40 | 43 | 40 | 40 | 42 | 40 | 78 | Yes | | | | | 4.25 | 0.0019 | 0.59 | EAANQ | | | | |
| 6 | Mucor circinelloides f. lusitanicus CBS 277.49 | OAD01005.1 | 27 | 76 | 72 | 90 | 100 | 100 | 73 | 66 | 66 | 97 | 68 | 68 | 73 | 40 | 43 | 40 | 40 | 42 | 40 | 78 | Yes | | | | | 4.25 | 0.0019 | 0.59 | EAANQ | | | | |
| 7 | Rhizopus azygosporus | RCH82711.1 | 27 | 85 | 87 | 78 | 73 | 73 | 100 | 77 | 77 | 73 | 77 | 77 | 76 | 45 | 46 | 41 | 43 | 48 | 45 | 68 | Yes | | | | | 4.25 | 0.0019 | 0.59 | EAANQ | | | | |
| 8 | Rhizopus microsporus | ORE22231.1 | 27 | 75 | 77 | 69 | 66 | 66 | 77 | 100 | 100 | 66 | 99 | 99 | 69 | 46 | 45 | 46 | 42 | 45 | 42 | 66 | Yes | | | | | | | | | | | | |
| 9 | Rhizopus stolonifer | RCH89556.1 | 27 | 75 | 77 | 69 | 66 | 66 | 77 | 100 | 100 | 66 | 99 | 99 | 69 | 46 | 45 | 46 | 42 | 45 | 42 | 66 | Yes | | | | | | | | | | | | |
| 10 | Mucor ambiguus | GAN06585.1 | 27 | 75 | 72 | 90 | 97 | 97 | 73 | 66 | 66 | 100 | 67 | 67 | 71 | 40 | 43 | 40 | 40 | 42 | 39 | 77 | Yes | | | | | 4.25 | 0.0019 | 0.59 | EAANQ | | | | |
| 11 | Rhizopus microsporus ATCC 52813 | XP_023462404.1 | 26 | 74 | 77 | 69 | 68 | 68 | 77 | 99 | 99 | 67 | 100 | 100 | 69 | 44 | 45 | 45 | 41 | 44 | 42 | 67 | Yes | | | | | | | | | | | | |
| 12 | Rhizopus microsporus var. microsporus | ORE05386.1 | 26 | 74 | 77 | 69 | 68 | 68 | 77 | 99 | 99 | 67 | 100 | 100 | 69 | 44 | 45 | 45 | 41 | 44 | 42 | 67 | Yes | | | | | | | | | | | | |
| 13 | Phycomyces blakesleeanus NRRL 1555(-) | XP_018295414.1 | 26 | 80 | 77 | 73 | 73 | 73 | 76 | 69 | 69 | 71 | 69 | 69 | 100 | 38 | 41 | 42 | 39 | 42 | 36 | 68 | | Yes | | | | 4.25 | 0.0019 | 0.59 | EAANQ | | | | |
| 14 | Absidia glauca | SAL96273.1 | 24 | 45 | 45 | 43 | 40 | 40 | 45 | 46 | 46 | 40 | 44 | 44 | 38 | 100 | 69 | 47 | 41 | 42 | 44 | 40 | Yes | | | | | | | | | | | | |
| 15 | Absidia repens | ORZ12009.1 | 23 | 45 | 48 | 45 | 43 | 43 | 46 | 45 | 45 | 43 | 45 | 45 | 41 | 69 | 100 | 43 | 38 | 39 | 44 | 41 | Yes | | | | Yes | | | | | | | | |
| 16 | Syncephalastrum racemosum | ORY99019.1 | 21 | 46 | 44 | 44 | 40 | 40 | 41 | 46 | 46 | 40 | 45 | 45 | 42 | 47 | 43 | 100 | 44 | 46 | 44 | 41 | | Yes | | | | | | | | | | | |
| 17 | Hesseltinella vesiculosa | ORX62783.1 | 21 | 45 | 44 | 41 | 40 | 40 | 43 | 42 | 42 | 40 | 41 | 41 | 39 | 41 | 38 | 44 | 100 | 56 | 51 | 36 | Yes | | | Yes | | | | | | | | | |
| 18 | Lichtheimia corymbifera JMRC:FSU:9682 | CDH57241.1 | 20 | 47 | 48 | 45 | 42 | 42 | 48 | 45 | 45 | 42 | 44 | 44 | 42 | 42 | 39 | 46 | 56 | 100 | 47 | 42 | | Yes | | | | 4.88/ 4.74 | 0.0009/ 0.0014 | 0.59/ 0.59 | GAAAF /EATRQ | | | | |
| 19 | Lichtheimia ramosa | CDS03748.1 | 17 | 43 | 44 | 42 | 40 | 40 | 45 | 42 | 42 | 39 | 42 | 42 | 36 | 44 | 44 | 44 | 51 | 47 | 100 | 41 | Yes | | Yes | | | 4.88/ 4.74 | | | | | | | |
| 20 | Parasitella parasitica | CEP13132.1 | 16 | 73 | 71 | 82 | 78 | 78 | 68 | 66 | 66 | 77 | 67 | 67 | 68 | 40 | 41 | 41 | 36 | 42 | 41 | 100 | Yes | | | | | 4.25 | 0.0019 | 0.59 | EAANQ | | | | |

**Note:**

1. Protein sequences were aligned using MUSCLE in CLUSTAL format and % sequence identify matrix was generated.
2. Protein sequences were scanned for individual motifs using FIMO (Find Individual Motif Occurrences): Version 5.1.1 tool (http://meme-suite.org).
3. The p-value (threshold p=0.001) of a motif occurrence is defined as the probability of a random sequence of the same length as the motif matching that position of the sequence with as good or better a score.
4. The score (Threshold +0.001) for the match of a position in a sequence to a motif is computed by summing the appropriate entries from each column of the position-dependent scoring matrix that represents the motif.
5. The q-value of a motif occurrence is defined as the false discovery rate if the occurrence is accepted as significant.
